# Driving Forces underlying Selectivity Filter Gating in the MthK Potassium Channel

**DOI:** 10.1101/2022.03.29.486236

**Authors:** Wojciech Kopec, Andrew S. Thomson, Bert L. de Groot, Brad S. Rothberg

## Abstract

K^+^ channel activity can be limited by C-type inactivation, which is likely initiated in part by dissociation of K^+^ ions from the selectivity filter, and modulated by side chains surrounding the selectivity filter. Whereas crystallographic and computational studies have linked inactivation to a ‘collapsed’ selectivity filter conformation in the KcsA channel, the structural basis for selectivity filter gating in other K^+^ channels has been less clear. Here, we combined electrophysiological recordings with molecular dynamics based, *in silico* electrophysiology simulations, to study selectivity filter gating in the model potassium channel MthK and its V55E mutant (analogous to KcsA E71) in the pore-helix. Experimentally, we find that MthK V55E has a lower open probability than the WT channel, due to decreased stability of the open state, as well as a lower unitary conductance. Simulations account for both aspects of these observations on the atomistic scale, showing that ion permeation in V55E is altered by two distinct orientations of the E55 side chain. In the ‘vertical’ orientation of E55, in which E55 forms a hydrogen bond with D64 (as observed with KcsA WT channels), the filter displays reduced conductance compared to MthK WT. In contrast, with ‘horizontal’ orientation, K^+^ conductance is closer to MthK WT; however the selectivity filter stability in the conducting conformation is lowered, and the filter more readily transitions to the inactivated conformation. Surprisingly, these transitions of MthK WT and V55E channels to the non-conducting (inactivated) state observed in simulations are associated with a widening selectivity filter, unlike its narrowing seen in KcsA, and reminisce the recent structures of stably-inactivated, voltage-gated potassium channels: *Shaker* W434F and Kv1.2 W362F mutants, as well as WT Kv1.3 channels.

## INTRODUCTION

Potassium (K^+^) channels are critically important in the control of many cellular functions, including action potential firing, neurotransmitter and hormone release, and muscle contraction [1]. A great deal of effort has been focused on understanding the molecular basis of K^+^ permeation and channel gating, and it is now clear that K^+^ channels gate at multiple structural loci [2]. For example, in voltage-gated K^+^ channels as well as inwardly-rectifying K^+^ channels, the pore-lining helices from each subunit can form a constriction (activation gate) at the cytosolic entrance to the pore (S6 gating) [3–5], whereas some these and other K^+^ channels can also undergo conformational changes near the selectivity filter, at the extracellular end of the pore (SF gating), to gate ion permeation (**Fig. 1 A, B**) [6,7]. A combination of structural and functional studies has revealed that the mechanisms underlying each of these gating processes are unlikely to arise from a single blueprint, and that diversity among K^+^ channels is likely to have yielded several complementary mechanisms to underlie gating that may be coupled to transmembrane voltage, ligands, or other stimuli [8–11].

**Figure 1.**
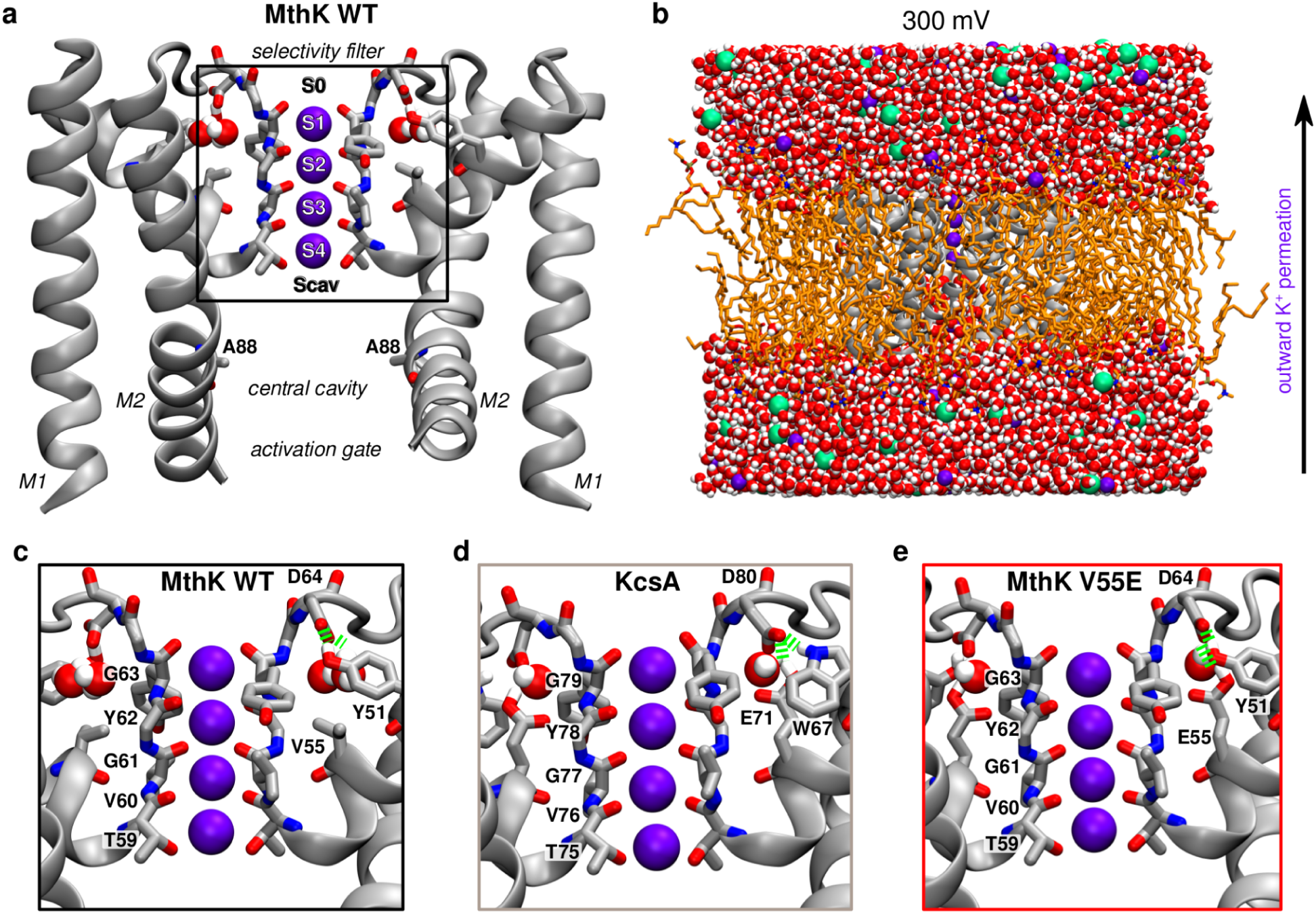
Basic architecture of a potassium channel. A, Crystal structure of MthK in an open state (PDB ID: 3LDC, only two diagonally opposite subunits of a tetramer are shown), with highlighted important elements - the selectivity filter in its conducting conformation consists of 4 main potassium binding sites S1-S4 and additional binding sites S0 and Scav. The central cavity, typically filled with water molecules, is located below the filter and above the activation gate. B, All-atom system used in MD simulations. The channel is shown in grey, lipids (POPC) in orange, water as red-white spheres, potassium ions as purple spheres and chloride ions as green spheres. Transmembrane voltage of 300 mV was used in simulations, resulting in an outward permeation of potassium ions through the channel as indicated by the arrow. C-E, Closer view on an interaction network behind the selectivity filter. In wild-type MthK, D64 makes hydrogen bonds (green springs) with Y51 and two crystallographic water molecules (C). In KcsA (PDB ID: 1K4C, D) a residue corresponding to Y51 is W67. Additionally protonated glutamate (E71, V55 in MthK) takes part in the same interaction network, with only one water molecule. The model of MthK V55E (E) is based on MthK WT, with V55 replaced with protonated E71 from KcsA and one water molecule.

It was observed previously that the prokaryotic K^+^ channel MthK exhibits SF gating that shares many properties in common with C-type inactivation of Kv channels. Specifically, channel opening is greatly decreased by increasing depolarization and decreasing extracellular potassium concentration ([K^+^]), and the gating process shows a selective sensitivity to cationic channel blockers, similar to that observed with the modulatory effect of quaternary ammonium derivatives on C-type inactivation in the *Shaker* K^+^ channel [9,12–16]. Interestingly, pore-helix glutamate that is a key determinant of inactivation in KcsA (E71) does not play a role in MthK gating; instead, this position contains a hydrophobic valine side chain (V55, **Fig 1. C, D**), identical to *Shaker* (V438), as well as mammalian Kv1.2 (V366) and Kv1.3 (V440) channels. This implies that the molecular driving forces and structural changes underlying SF gating among these channels may be different from one another.

To better understand the molecular basis for SF gating in MthK, we examined the functional consequences of substituting V55 in MthK with (protonated) glutamate (**Fig 1. E**) by electrophysiology, and analyzed potential underlying molecular effects by *in silico* electrophysiology molecular simulation. We observe that the V55E mutation yields a gating phenotype with markedly reduced open state stability, manifested as decreased channel open probability and decreased open times over a wide range of voltages. Further, the V55E mutation also decreases the unitary conductance of MthK channels.

On the molecular level, simulations reveal two distinct orientations of the E55 side chain (‘vertical’ and ‘horizontal’) that differentially affect the SF properties. The ‘vertical’ orientation is reminiscent of E71 seen in structures of KcsA (**Fig 1. D**), where it is hydrogen-bonded with an aspartate behind the SF (D80 in KcsA; D64 in MthK). In this orientation, MD simulations predict a low conductance of MthK V55E. In the second, ‘horizontal’ orientation, that is with the E55 side chain oriented perpendicular to the membrane, K^+^ conductance is closer to MthK WT, but longer simulations show that this orientation promotes SF entering into the non-conducting (inactivated) state. In MD simulations, SFs of both WT and V55E channels undergo gating transitions that share similarities with the C-type inactivated conformations observed in stably-inactivated mutants of *Shaker* and Kv1.2 channels, and in WT Kv1.3 channels. In a full accord with electrophysiology, V55E SF shows in simulations a larger degree of instability than WT, which arises from the entry of additional water molecules behind the SF and K^+^ ion unbinding from the top of the SF. Taken together, these data suggest a mechanism for SF gating (inactivation) in MthK channels, and highlight the critical role for interactions between the SF and pore-helix in stabilizing the K^+^ ion conduction pathway. Finally, we highlight several important differences between two popular molecular force fields (CHARMM36m and AMBER14sb) used in MD simulations of potassium channels.

## RESULTS

### The V55E mutation decreases stability of the open state

Mutation of E71 to alanine, in the pore-helix of KcsA, was observed to abrogate a key interaction between the side chain carboxylates of E71 and D80; elimination of this interaction apparently results in stabilization of the open state and loss of inactivation in KcsA channels [6]. It was observed previously that MthK channels can also exhibit an inactivation-like phenomenon, in which the open state is stabilized by occupancy of a K^+^-selective site at the external end of the SF [9,12].

To determine whether MthK inactivation and KcsA inactivation might arise through similar molecular interactions, we examined functional consequences of substituting MthK V55 (at the position analogous to KcsA E71) with glutamate. We observe that with symmetrical 200 mM K^+^ solutions, MthK WT channels, activated by 100 µM Cd^2+^ at the cytosolic side of the channel, gate with high open probability (Popen) of >0.9 at voltage ranging from −150 to +140 mV (**Fig. 2**) [12,17,18]. At strongly depolarized voltages (>140 mV), Popen decreases sharply with increasing depolarization, with a voltage-dependence of e-fold per 14 mV and a V_1/2_ for this decrease at 190 ± 1.5 mV (7). This robust decrease in Popen is enhanced by decreasing external [K^+^] (K^+^_ext_), such that a decrease from 200 mM to 5 mM K^+^_ext_ shifts the V_1/2_ leftward from 190 to 95 ± 2.6 mV (**Fig. 2 C**) [12]. These gating effects are described by a model in which voltage-dependent outward K^+^ ion movement, followed by dissociation of a K^+^ ion from the external end of the SF, leads to a conformational change within the SF that consists of an off-axis rotation of a carbonyl group (at V60) that normally lines the selectivity filter [9,12]. This carbonyl rotation can halt conduction and thus gate the channel [12].

**Figure 2.**
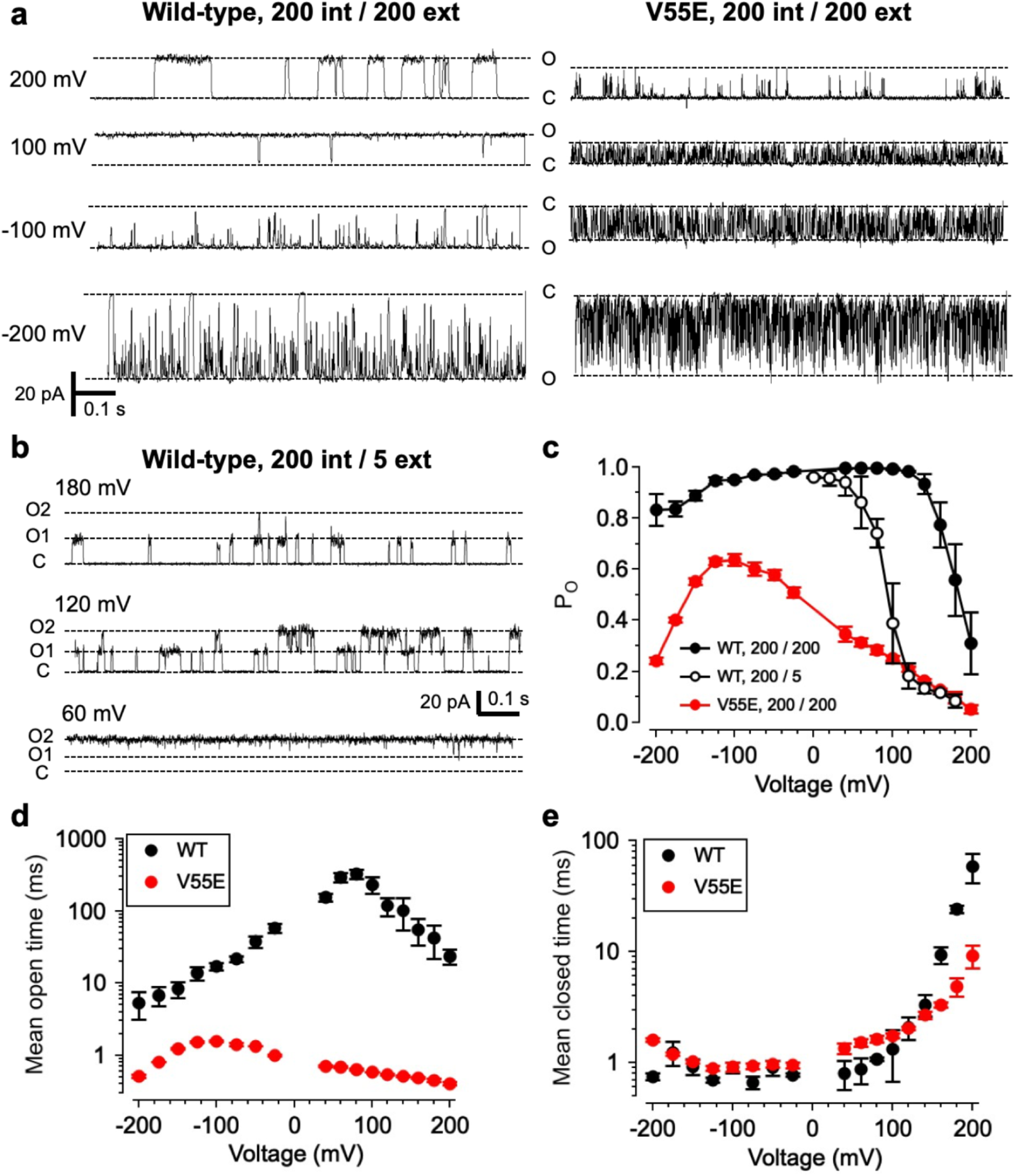
Effects of V55E mutation on MthK channel gating. A, Representative single-channel current through MthK wild-type and V55E channels over a range of voltages. For these recordings, solutions at both sides of the membrane contained 200 mM KCl. “O” and “C” indicate open and closed channel current levels, respectively. Strong depolarization (200 mV) leads to decreased open probability (Po) for wild-type channels, and V55E channels consistently display briefer openings than wild-type at all tested voltages. B, current through MthK wild-type channels with 5 mM external K^+^. This bilayer contained two active channels, indicated by open levels “O1” and “O2”. Lower external [K^+^] enhances inactivation at mildly depolarized voltages. C, Po vs. voltage for wild-type (WT) channels with 200 mM (black circles, n=5 bilayers) or 5 mM external K^+^ (open circles, n=3 bilayers), compared with V55E with 200 mM external K^+^ (n=5 bilayers). D, MthK WT channel open times were voltage dependent, with a maximal mean open time of 1200 +/- 120 ms at 60 mV; in contrast V55E were much briefer, with the mean open time not exceeding 6.7 +/- 0.2 ms over a 400 mV range. E, WT and V55E closed times were similar over a wide range of voltages, consistent with the idea that the major effect of the V55E mutation was to destabilize the open/conducting state relative to WT. [K^+^]_ext_ = 200 mM (WT n=5; V55E n=5).

In contrast, with symmetrical 200 mM K^+^ solutions, the MthK V55E mutant reaches a maximum Popen of 0.64 ± 0.02 at −100 mV, and Popen decreases with either depolarization or stronger hyperpolarization, reaching a value of 0.054 ± 0.02 at 200 mV and 0.24 ± 0.01 at −200 mV (**Fig. 2 C**). Over the entire 400 mV voltage range examined in these experiments (from 200 to +200 mV), MthK V55E channels exhibited lower Popen than MthK WT channels recorded under the same conditions.

The single-channel currents in **Fig. 2 A** suggested that the basis for the decreased Popen in MthK V55E channels may be a decrease in mean open time compared to WT. Consistent with this observation, **Fig. 2 D** illustrates that the mean open time of V55E channels is much briefer than WT over voltages ranging from −200 to +200 mV. Mean open times of V55E channels were 10- to over 100-fold briefer than WT, with a maximal mean open time of 1.6 ± 0.03 ms at −100 mV and 0.41 ± 0.02 ms at 200 mV for V55E channels. In contrast, the mean open time of MthK WT channels reached a maximum of 330 ± 46 ms at +80 mV, and decreased to 23.6 ± 5.3 ms at 200 mV.

Although the durations of openings were markedly different between V55E and WT channels, closed times of V55E channels were very similar to WT over a wide range of voltages up to 140 mV (**Fig. 2 E**). Interestingly, although V55E channels are characterized by lower Po than WT channels over a wide range of voltages, WT channels show an increase toward longer mean closed times than V55E at voltages greater than 140 mV (at 200 mV, mean closed time of WT, 58.2 ± 17 ms; V55E, 9.11 ± 2.1 ms). This is consistent with the idea that destabilization of the open state in V55E channels, rather than increased stabilization of the closed state, is the critical determinant of the gating phenotype in V55E channels. Together, these observations indicate that the V55E mutation leads to decreased lifetime, and thus decreased stability of the open (conducting) state.

### The V55E mutation decreases unitary conductance

In single-channel recordings, one can observe that in addition to decreased open times, V55E channel openings frequently do not reach the same amplitude as WT openings (**Fig. 2 A**). Measuring the amplitudes of unitary currents for V55E channels is made complicated by the observation that V55E openings are very brief (**Fig. 2 D; Fig. 3**), and amplitudes of brief events are truncated by the effects of filtering, such that with the 1 kHz low-pass filtering for recordings in these studies (resulting in a dead-time of 0.179 ms; see Methods), the amplitudes of openings with durations less than 0.716 ms (dead-time x 4) will be truncated [19]. In addition, channel openings can occur in bursts containing brief closings, and very brief closings that fail to cross the threshold could also affect estimates of unitary current amplitude.

**Figure 3.**
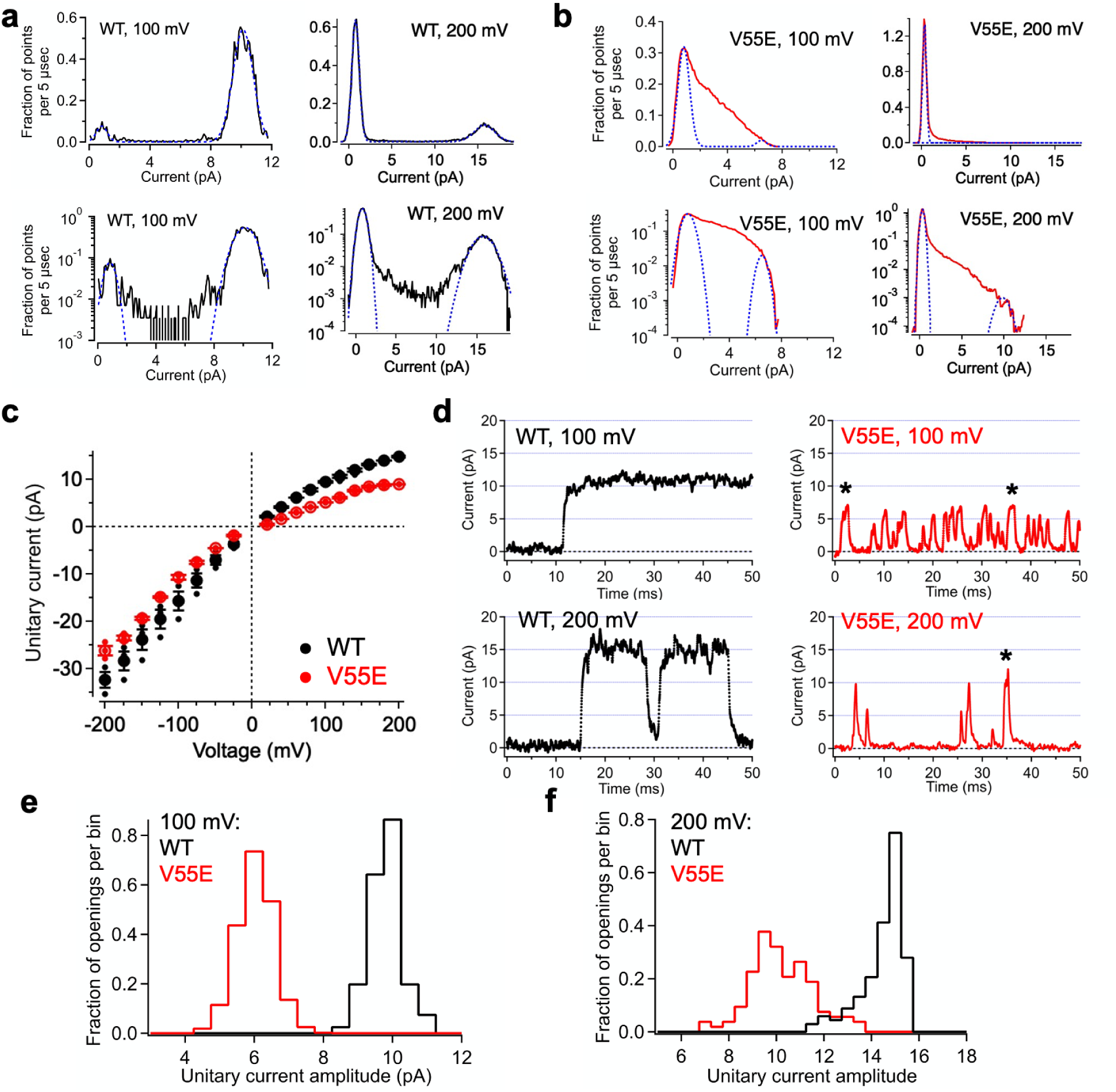
Effects of V55E mutation on MthK channel conductance. A, B, Representative all-points histograms from a WT and a V55E channel, respectively, constructed from selected data segments at 100 and 200 mV. Peaks correspond to open and closed current levels. These were fitted with two Gaussian components (blue dashed lines), and the difference between the means yields an estimate of the current amplitude. Upper panels show histograms on a linear scale; the same histograms are shown on a logarithmic scale on the corresponding lower panels. C, Unitary current vs. voltage for WT and V55E channels, determined from all-points histograms at voltages ranging from −200 to 200 mV. Data points represent means ± SEM (smaller points represent data from individual experiments). D, Representative single-channel openings from wild-type and V55E channels at 100 mV (top panels) and 200 mV (lower panels). V55E openings with durations >1.074 ms are indicated by an asterisk. E, Histograms of amplitudes of single-channel openings with durations >1.074 ms at 100 mV, each normalized to an area of 1.0 (WT, 81 openings; V55E, 963 openings). Mean unitary currents for these openings were: WT 9.6 ± 0.5 pA; V55E 5.8 ± 0.5 pA (mean ± S.D.). F, Histograms as in panel E from openings at 200 mV (WT, 136 openings; V55E, 106 openings). Mean unitary currents for these openings were: WT 14.2 ± 0.9 pA; V55E 10.0 ± 1.3 pA (mean ± S.D.).

To gain insight toward whether the V55E mutation might reduce the unitary conductance of the channel, we quantified unitary current amplitudes over a range of voltages by constructing all-points histograms, and determined the difference between mean open and closed (baseline) current levels based on the peaks of these histograms. Representative histograms of WT and V55E currents at 100 and 200 mV are shown in **Fig. 3 A, B**. For WT channels, all-points histograms appeared as bimodal distributions, corresponding to the closed and open current levels for these channels (**Fig 3 C**). Each peak was fitted with a Gaussian distribution (*blue dashed curves*) to estimate the mean open and closed current levels, and unitary current was determined as the difference between the levels. So, for example, the WT open channel current amplitude at 200 mV from the representative all-points histogram in **Fig 3 A** was 14.9 pA (15.7 minus 0.78 pA), and the mean WT amplitude at 200 mV was 14.78 ± 0.17 pA (n=3).

For V55E channels, unitary current measurements by the all-points histogram approach (**Fig 3 B**) were complicated by the skewed distribution of points in the intermediate current range between open and closed. The shapes of the V55E all-points histograms can be attributed to the overrepresented time spent in transitions between opening and closing in V55E channels; because V55E openings were so brief, a large fraction of the digitized signal comprised transitions between open and closed levels. In addition, the low open probability of these data resulted in very small peaks for the open levels in most of the data sets. To better resolve both the maximum and minimum current levels, we plotted the all-points histograms on semi-logarithmic coordinates (lower panels in **Fig 3 A, B**). Using semi-logarithmic coordinates, one can observe the rapid fall-off at the ends of the histograms; these were well-described by the tails of the Gaussian functions (blue dashed curves) to enable estimates of the open and closed levels. Thus, for example, although the open probability for V55E channels at 200 mV was low (∼0.05), **Fig 3 B** (*lower right panel*) shows that the maximum open current level at 200 mV could be resolved as 9.8 pA, yielding an estimate of open amplitude as 9.5 pA (9.8 minus 0.34 pA); the mean V55E amplitude at 200 mV for channels in this analysis was 8.98 ± 0.040 pA (n=3), which was significantly lower than the WT current amplitude (14.8 ± 0.17 pA) measured in the same conditions (t-test; p<0.001).

As an additional check on our estimates of open current levels from WT and V55E channels, we measured the amplitudes of individual openings in conjunction with measurement of dwell times (see Methods). As noted above, amplitudes of brief openings with durations < 0.716 ms are truncated due to the effects of low-pass filtering [19]. Although in principle, openings with durations >0.716 ms should be long enough to reach the “full” open current level, openings very close to this threshold would yield amplitude estimates based on only a few digitized points, and would thus be vulnerable to error due to the presence of noise (up to ± 0.4 pA). We therefore used a threshold of 1.074 ms (dead-time x 6) for openings included in this analysis. Representative openings meeting the 1.074 ms criterion are shown in **Fig. 3 D** (long duration V55E openings are indicated by asterisk). Histograms constructed from WT and V55E unitary current measurements at 100 and 200 mV (**Fig. 3 E, F**) illustrate that long duration V55E openings have significantly lower amplitudes than WT amplitudes. For openings at 100 mV, the mean amplitude for WT was 9.6 ± 0.5 pA, while V55E was 5.8 ± 0.5 pA; at 200 mV, WT openings were 14.2 ± 0.9 pA, while V55E openings were 10.0 ± 1.3 pA (given as mean ± S.D.). These measurements were consistent with those obtained using all-points histograms, together supporting the idea that the V55E mutation underlies altered channel function in terms of both reduced stability of the open conformation and ion conduction.

### Shorter in silico electrophysiology simulations - AMBER14sb force field

To probe the effects of the V55E mutation in the MthK channel on an atomistic scale, we used our previous computational protocol to monitor changes in outward current, caused by motions of the inner helices (the activation gate, **Fig. 4 A**) [21]. Initially, we used the AMBER14sb force field (referred to as AMBER, see Methods for details). When compared to the WT channel, simulated currents at 300 mV in MthK V55E display a similar overall behavior, with an initial increase in magnitude upon the activation gate opening (measured as an average distance between CA of A88 residues from oppositely oriented monomers of MthK, **Fig. 4 B**), that is between 1.4 - 1.6 nm, from the initial value of 2-3 pA. Around the opening of 1.6 nm, the current reaches its maximum of ∼16 pA in WT and ∼10 pA in V55E. Further activation gate opening (above the value of 1.6 nm) leads to a current decrease, back to 2-3 pA in both channels. In WT, we have previously traced this behavior to the variations in the width of the S4 binding site, formed exclusively by T59 (**Fig. 4 A)**. Indeed, when T59 CA-CA distance (i.e. the width of the S4 ion binding site) is used as an opening coordinate, currents in V55E show a similar trend upon S4 opening as in WT (**Fig. 4 C**). In contrast however, the maximum outward current in V55E at 300 mV (∼10 pA) is markedly lower than the WT maximum current (∼16 pA), suggesting that an additional energetic barrier for ion permeation (apart from the one imposed by T59) exists somewhere along the permeation pathway.

**Figure 4.**
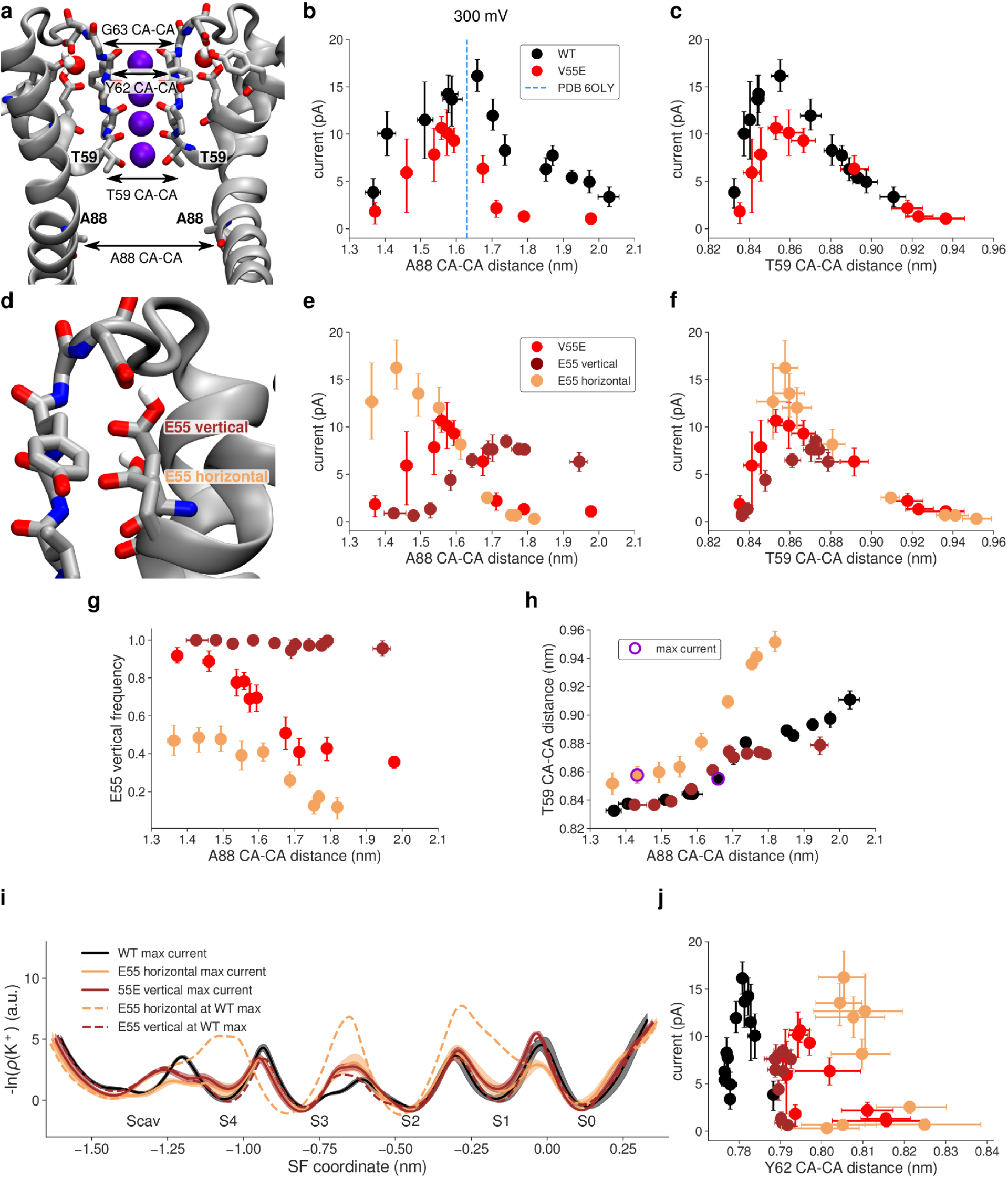
Outward currents in MthK V55E in MD simulations with the AMBER force field. A, Starting structure of MthK V55E and the definition of distances used as opening coordinates (A88 CA-CA distance: activation gate; T59 CA-CA distance: S4 width). B, Outward current through MthK WT (black traces) and MthK V55E (red traces) as a function of the activation gate opening. Blue vertical line marks the largest opening level seen experimentally. C, Same as B, but with the S4 width used as the opening coordinate. D, Two rotameric states (orientations) of E55 observed in MD simulations. E, Outward current in ‘E55 vertical’ (brown traces) and ‘E55 horizontal’ (light orange traces) simulation sets in comparison to current in V55E (red traces), as a function of the activation gate opening. F, Same as E, but with the S4 width used as the opening coordinate. G, Frequency of the ‘vertical’ rotameric state of E55 as a function of the activation gate opening. H, Correlation between S4 width and the activation gate opening in ‘E55 vertical’ and ‘E55 horizontal’ in comparison to WT. Points for which the maximal current of ∼16 pA was recorded are indicated by a violet ring. I, Negative logarithmic densities (‘free energy profiles’) of K^+^ ions in the SF. Minima correspond to stable ion binding sites and maxima to free energy barriers between them. J, Currents in all systems with the Y62 (forming S1 and S0 sites) CA-CA distance used as an opening coordinate. On all panels, error bars represent 95% confidence intervals.

In simulations of MthK V55E, we observed that the E55 side chain adopts two distinct rotameric states - one that is seen in crystal structures of KcsA, where the (protonated) side chain of E55 (E71 in KcsA) is hydrogen-bonded with the side chain of D64 (D80 in KcsA), behind the SF (‘vertical’ orientation of E55, **Fig. 4 D**). In the second orientation, this hydrogen bond is broken, and instead the side chain of E55 adopts a more in-plane orientation with respect to the membrane (‘horizontal’ orientation of E55). We suspected that this rotameric transition might have a large influence on the conducting properties of the SF. For a clearer picture, we performed separate simulations with the E55 side chain weakly restrained to one of the two orientations (simulation sets ‘E55 vertical’ and ‘E55 horizontal’). These simulations show that the orientation of the E55 side chain has a major effect on the outward current, and on its dependence on both gate and S4 opening (**Fig. 4 E, F**).

In ‘E55 vertical’ simulations, the currents are markedly lower than in the non-restrained case, at low and moderate activation gate openings, and only at larger openings reach the level of 7-8 pA (**Fig. 4 E**, brown traces). Furthermore, the current does not decrease at very large openings. The reason for that is clear when the S4 width is used as the opening coordinate (**Fig 4 F**, brown traces): in the ‘E55 vertical’ simulations, the S4 site does not get wider than approximately 0.88 nm, showing that the activation gate S4 coupling is restricted when the E55 side chain adopts the ‘vertical’ orientation, in contrast to non-restrained simulations of V55E (and WT, see **Fig. 4 C**). In ‘E55 horizontal’ simulations, the current reaches its maximum value at small activation gate openings of ∼1.43 nm, in contrast to 1.6-1.65 nm in the non-restrained case (and WT) (**Fig. 4 E**, light orange traces). Interestingly, the S4 width in ‘E55 horizontal’ is identical to MthK WT, at openings corresponding to the maximum current. In other words, the ‘horizontal’ orientation of the E55 side chain affects the activation gate - S4 coupling: the S4 binding site is able to open wider at smaller activation gate openings. Note however, that the restraints used were sufficiently mild, that there is still a significant portion of ‘vertically’-oriented E55 side chains in these simulations (**Fig 4 G)**. The maximum current in the ‘E55 horizontal’ simulations reaches the value of MthK WT, further demonstrating that MthK WT and V55E can, in principle, conduct ions at a similar rate. Overall, currents in non-restrained MthK V55E, are affected not only by the activation gate opening (as in WT), but additionally also by the rotameric state of the E55 side chain (**Fig. 4 E,** red traces).

We next sought to determine why in these simulations are V55E currents reduced compared to MthK WT or ‘E55 horizontal’ simulations, even though in all these cases the optimal width of the S4 site is reached (∼0.86 nm, **Fig. 4 F, H**). To gain additional insights, we calculated negative logarithms of potassium densities in the SF that might be treated as approximate, one dimensional free energy profiles for ion permeation (**Fig. 4 I**). It is important to note that such profiles do not reflect the underlying multi-ion permeation process, including permeation barriers. They indicate the regions in the SF where ion binding is favored and disfavored.

A comparison between profiles for MthK WT vs. MthK V55E (black curve vs. solid brown and light orange curves in **Fig. 4 I**) reveals that the barriers are reduced at the entry to the SF, for both orientations of the E55 side chain (i.e. between Scav and S4, and between S4 and S3). However, in ‘E55 vertical’ simulations, the barrier between S1 and S0 is increased compared to WT (both brown curves). Consequently, the width of the SF between S1 and S0 (i.e. the CA-CA distance of Y62) does not show any variation in ‘E55 vertical’ simulations, being restricted to the value of ∼0.79 nm, in stark contrast to both MthK WT and ‘E55 horizontal’ simulations (**Fig. 4 J**). Interestingly, this value lies between those observed in MthK WT and ‘E55 horizontal’ simulations suggesting that this particular value leads to a very strong binding of a K^+^ ions to S2. Indeed in ‘E55 vertical’ simulations the occupancy of S2 by K^+^ ions is >0.95, compared to ∼0.82 in WT.

In ‘V55E horizontal’ simulations, at an opening that shows the maximum current, the free energy profile is quite flat with low barriers, consistent with the observed high current (**Fig. 4 I**, light orange curve). The profile at a larger opening (dashed curve) reveals the effect of the ‘horizontal’ orientation of E55 on S4 in the AMBER force field: the S4 binding site is compromised (a free energy minimum is replaced by a maximum), which likely results in to two other high barriers, namely between the S3 and S2 as well as S2 and S1 ion binding sites, drastically decreasing the current at larger openings in ‘E55 horizontal’ simulations.

### Shorter in silico electrophysiology - CHARMM36m force field

Subsequently, we simulated the same systems with the CHARMM36m force field (referred to as CHARMM, see Methods for details), to see whether the previous observations are force field dependent. In our previous study of MthK WT, both force fields showed the same trend in the current dependence on the lower gate opening, as well as very similar maximum currents [21]. Overall, simulations of V55E done with the CHARMM force field show little currents (up to 1.5 pA, i.e. more than an order of magnitude smaller than in WT) at all levels of the lower gate opening (**Fig. 5 A**). The S4 does however become wider in V55E, similarly to WT (**Fig. 5 B**), suggesting that a large free energy barrier for ion permeation exists somewhere else in the SF. In contrast to AMBER simulations, where we observed two rotamers of the E55 side chain, in CHARMM the E55 side chain adopts almost exclusively the ‘vertical’ orientation (**Fig. 5 C,** red traces). Initial tries of inducing the ‘horizontal’ orientations were unsuccessful; we then used some of the ‘horizontal’ orientations seen in AMBER for further simulations with the CHARMM force field and imposing strong position restraints (**Fig. 5 C**, light orange traces). For completeness, we also simulated a system with distance restraints present in the ‘vertical’ orientation (**Fig. 5 C**, brown traces). This simulation set, ‘E55 vertical’, behaves almost identical as the unrestrained case, indicating that the application of restraints has negligible effects on the overall channel function.

**Figure 5.**
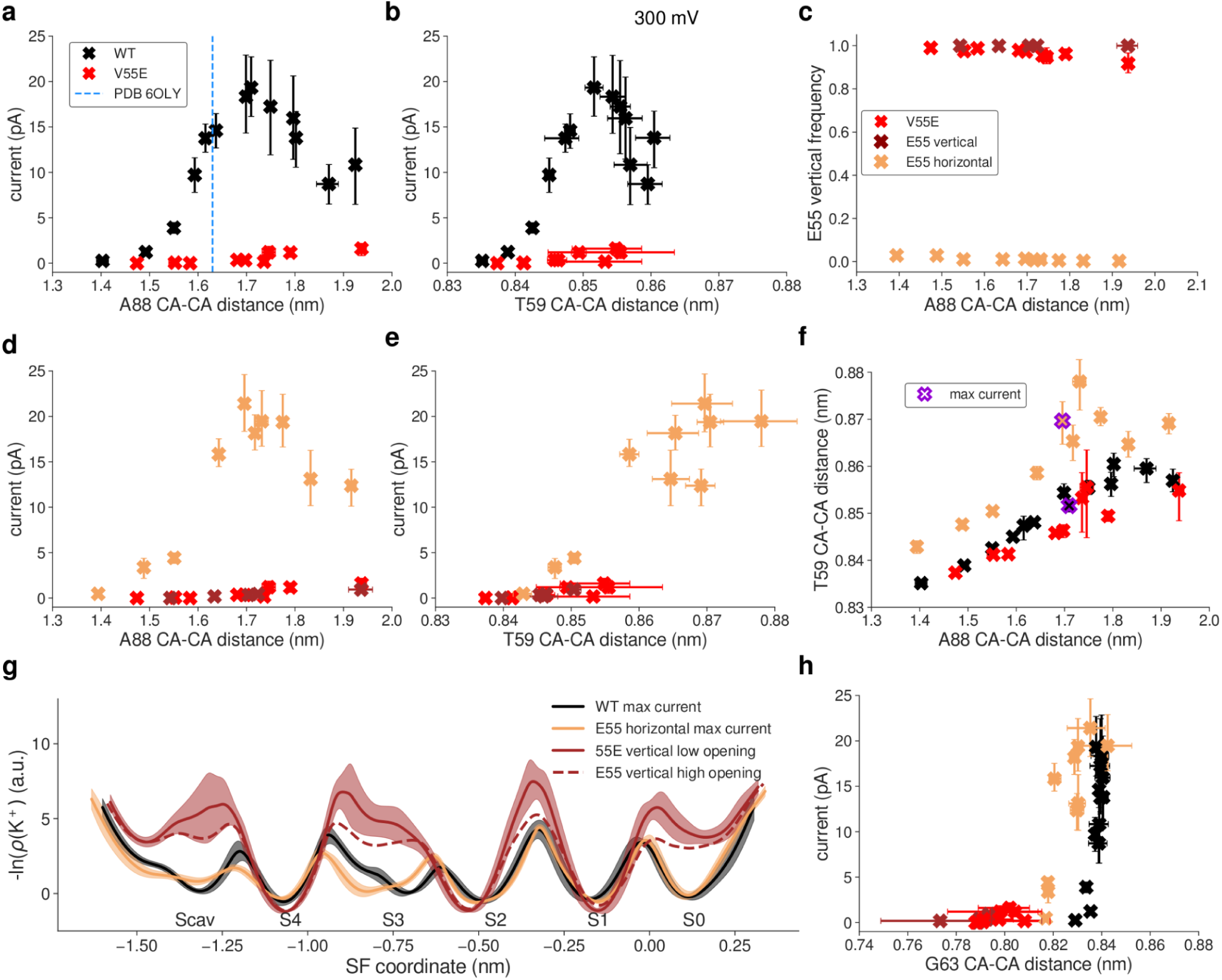
Outward currents in MthK V55E in MD simulations with the CHARMM force field. A, Outward current through MthK WT (black traces) and MthK V55E (red traces) as a function of the activation gate opening. Blue vertical line marks the largest opening level seen experimentally. B, Same as A, but with the S4 width used as the opening coordinate. C, Frequency of the ‘vertical’ rotameric state of E55 as a function of the activation gate opening. D, Outward current in ‘E55 vertical’ (brown traces) and ‘E55 horizontal’ (light orange traces) in comparison to current V55E (red traces), as a function of the activation gate opening. E, Same as D, but with the S4 width used as the opening coordinate. F, Correlation between S4 width and the activation gate opening in ‘E55 vertical’ and ‘E55 horizontal’ in comparison to WT. Points for which the maximal current of ∼20 pA was recorded are indicated by a violet cross. G, Negative logarithmic densities (‘free energy profiles’) of K ions in the SF. Minima correspond to stable ion binding sites and maxima to free energy barriers between them. H, Current in all systems with the G63 (forming S0 site) CA-CA distance used as an opening coordinate. On all panels, error bars represent 95% confidence intervals.

Similarly to simulations done with the AMBER force field, when the E55 side chain adopts the ‘horizontal’ orientation in the CHARMM force field, the channel is able to permeate K^+^ ions as efficiently as WT (**Fig. 5 D**). In CHARMM however, that happens at the same gate opening level as in WT, which somewhat unexpectedly corresponds to slightly larger openings of the S4 in ‘E55 horizontal’ (**Fig. 5 E, F**). These observations suggest that the free energy barrier imposed by T59 is not always the dominating one in CHARMM. Indeed, in the WT channel, there is a barrier of a similar height located between S2 and S1 binding sites (**Fig. 5 G**, black traces). This barrier becomes dominant in ‘E55 horizontal’ simulations, even though the barrier between S4 and S3 got lowered further, explaining the same maximal current in both WT and ‘E55 horizontal’ simulations.

The remaining question is, again, why is ion conduction in unrestrained V55E and ‘E55 vertical’ simulations so low (max. 1.5 pA in the CHARMM force field). The free energy profiles (**Fig. 5 G**, brown traces) show several very high barriers, as well as severely affected Scav and S0 ion binding sites. The lack of occupancy at the Scav binding site can be explained by a high ion occupancy at S4, which would electrostatically repel ions approaching Scav. However, the lack of the S0 binding site is more puzzling, also because it was preserved in AMBER simulations. The analysis of CA-CA distances of S0-forming G63 (**Fig. 5 H**) reveals that the ‘vertical’ orientation of the E55 side chain makes S0 too narrow, and in consequence it likely cannot bind K^+^ ions anymore. Only upon the E55 side chain rotation toward the ‘horizontal’ orientation, S0 becomes as wide as in MthK WT, which correlates with the increased current (**Fig. 5 H**, black and light orange traces).

Summarizing, CHARMM and AMBER force fields show marked differences in the exact behavior of the MthK V55E channel and in ion permeation through it. However, they are in a broad accord in three main points: i) ion permeation in MthK V55E depends not only on the coupling between the activation gate and S4 (as is does in WT channels), but also on the rotameric state of the E55 side chain; ii) the ‘vertical’ E55 orientation, i.e. hydrogen-bonded with both D64 and Y51 (**Fig. 1 E**), reduces channel conductance, by imposing additional free energy barriers for ion permeation in the SF; and iii) the ‘horizontal’ orientation of E55 increases currents in V55E to WT-like levels, by reducing these free energy barriers, especially in the upper part of the SF.

The reduced outward conductance of MthK V55E, as compared to MthK WT, predicted by both force fields, is in very good agreement with our electrophysiological unitary conductance measurements (**Fig. 3**). The major difference between the force fields in this set of simulations is the fact that in AMBER both ‘vertical’ and ‘horizontal’ orientations of the E55 side chain are frequently visited (**Fig. 4 G**), whereas CHARMM has a strong preference for the ‘vertical’ orientation (**Fig. 5 C)**. This, in turn, results in very different simulated outward currents when no restraints on the E55 side chain are used, because, as shown above, the orientation of this side chain is a major factor regulating ion permeation.

The origins of this distinct behavior, i.e. why the horizontal orientation is frequently visited in the AMBER force field, but not in CHARMM, are not obvious. It is unlikely however, that this difference arises from variations in the non-bonded interactions. Even though the partial charges and Lennard-Jones parameters are different between AMBER and CHARMM, the resulting interaction energies of the D64-E55 hydrogen bond (E55 in the ‘vertical’ orientation) are actually quite similar and favorable (**Supplementary Fig. 1 B, C**). However, we suspected that the energetics of the two dihedrals, that involve the hydrogen atom from the protonated E55 side chain, might also play a role (**Supplementary Fig. 1 D**). Indeed, these two dihedrals are low in energy (0-2 kJ/mol) in the ‘vertical’ orientation in the CHARMM force field (**Supplementary Fig. 1 E**). In contrast however, they are energetically less favorable in the same orientation in the AMBER force field, as one of the dihedrals has an energy of ∼17-18 kJ/mol (**Supplementary Fig. 1 F**). Only after the transition to the ‘horizontal’ orientation, the energy drops to values close to 0 kJ/mol, likely explaining the frequent transition from ‘vertical’ to ‘horizontal’ orientation in the AMBER force field (and lack of thereof in CHARMM). Further research is needed to unravel which of these force field parametrizations describe the E55 dynamics more realistically.

The orientational preference of E55 side chain in AMBER (but not in CHARMM) is also dependent on the activation gate opening (**Fig. 4 G**), which we attribute to the allosteric coupling between the gate and the SF, and the residues nearby [21,22]: a variation in the activation gate width affects the SF width as well, which in turn might impact the delicate balance between non-bonded interactions and torsional energetics, dictating the overall E55 side chain dynamics.

### Longer in silico electrophysiology simulations

Apart from faster protein motions occurring on the tens and hundreds of nanoseconds timescale studied above, we also investigated the SF gating (inactivation), i.e. conformational transitions of the SF, with MD simulations. Even though in reality, the timescales of such transitions are at least in the millisecond range [23], way above the standard MD range, they have nevertheless been observed in simulations in a few cases [24,25].

Structures of KcsA, stably-inactivated mutants of voltage-gated K^+^ channels (*Shaker* W434F and Kv1.2 W362F) and WT Kv1.3 channels revealed that C-type inactivation-related conformational changes at the SF can follow at least two distinct paths (**Fig. 6**) [26–29]: in KcsA the SF pinches (constricts) at the level of the ‘first’ glycine (G77, corresponding to G61 in MthK), which is accompanied by flipping of valine (V76 in KcsA, corresponding to V60 in MthK) carbonyl away from the SF axis. The SF also loses two K^+^ ions during inactivation, and has remaining ions bound at S1 and S4, whereas S3 is occupied by a water molecule, which presumably enhances the valine flipping. In voltage-gated channels the SF dilates (widens) at the level of the ‘second’ glycine (G446 in *Shaker,* G374 in Kv1.2, G448 in Kv1.3, corresponding to G63 in MthK). In this case, this SF dilation is accompanied by a breakage of hydrogen bonds between aspartate (D447 in *Shaker*, D375 in Kv1.2, D449 in Kv1.3, corresponding to D64 in MthK) and tyrosine/tryptophan (W434 in *Shaker*, W362 in Kv1.2, W436 in Kv1.3, corresponding to Y51 in MthK), which is replaced by a phenylalanine in stably-inactivated mutants, and flipping of aspartate side chains toward the extracellular side. In contrast, these hydrogen bonds are retained during SF inactivation in KcsA and the aspartate (D80) flipping has not been observed (**Fig. 6 B**). In *Shaker* and Kv1.2, two K^+^ ions remain bound to the SF in its inactivated conformation, but they occupy S3 and S4 ion binding sites (in contrast to inactivated KcsA), and S2 is occupied by a water molecule. In Kv1.3 there is a third K^+^ ion bound at the S1 binding site. Consequently, the valine flipping has not been observed in structures of inactivated Kv channels, because S3 is occupied by a K^+^ ion.

**Figure 6.**
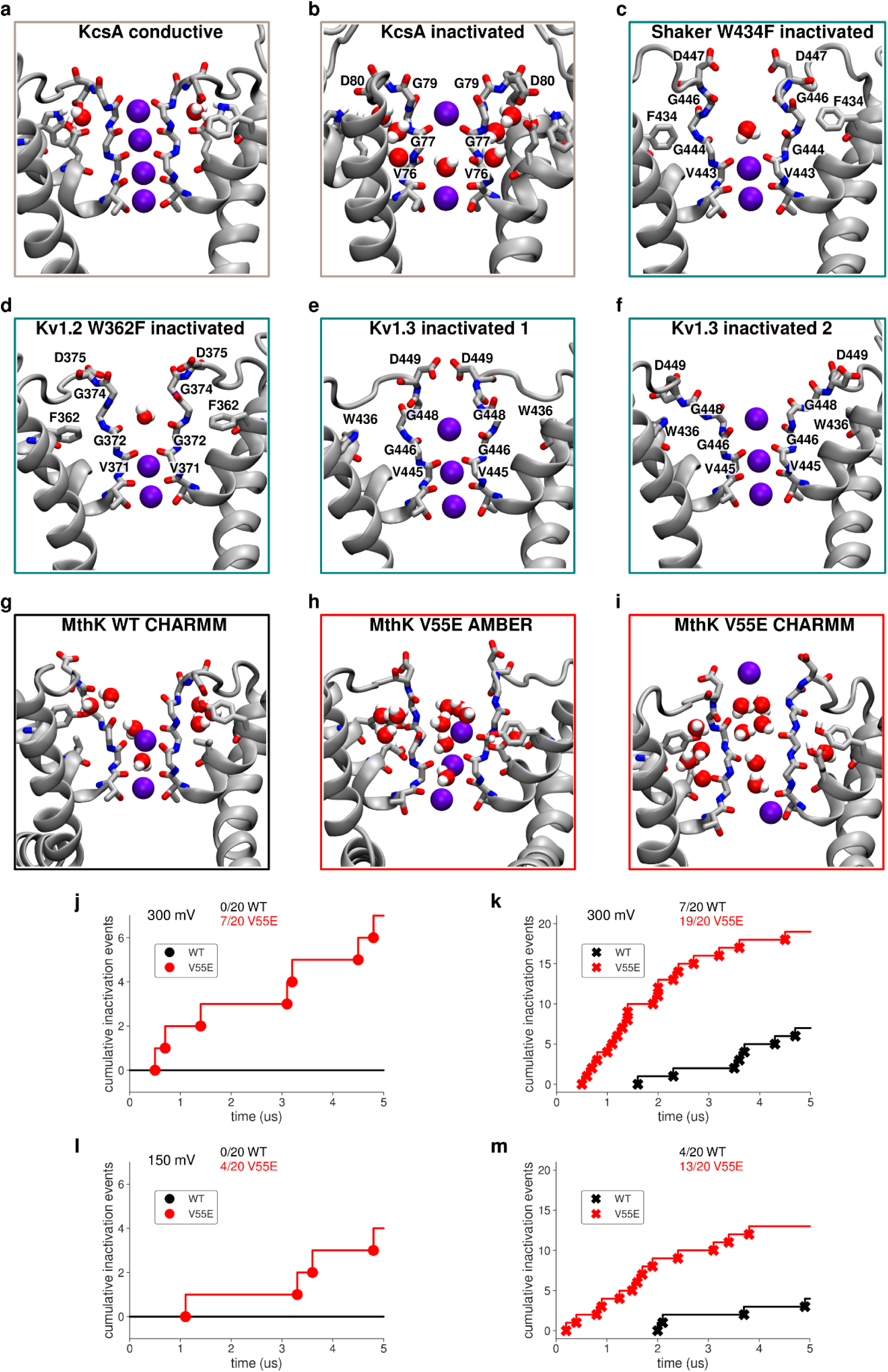
Conformations of inactivated filters in different K^+^ channels. A, Typical conducting SF conformation shown for reference (PDB ID: 1K4C). B, Inactivated (‘constricted’ or ‘pinched’) KcsA filter (PDB ID: 1K4D). The filter narrows at the level of the ‘first’ glycines (G77) and shows valine flipping at V76. C, Inactivated (‘dilated’) filter of *Shaker* W434F. D447 side chains flip toward the extracellular side, and the SF is widened/dilated at the level of the ‘second’ glycines (G446). D, Inactivated (‘dilated) filter of Kv1.2 W362F. Similarly to C, D375 side chains are flipped and the SF is widened. E, F, Two (1 and 2) inactivated conformations of Kv1.3. Again, D449 side chains are flipped and the SF is dilated. G-I, Examples of SF conformations observed in this work in long simulations of either MthK WT (G) or MthK V55E (H, I) at 300 mV. In all cases, flipping of aspartates (D64) is observed, together with widening at the level of the ‘second’ glycines (G63). Valine (V60) flipping is also observed as well. J-M, Cumulative inactivation events recorded in simulations with AMBER (J, L) and CHARMM (K, M) force fields at 300 and 150 mV, respectively. The events were identified based on distances between G63 CA atoms in oppositely oriented monomers.

Previous works on KcsA revealed that the SF pinching occurs in unbiased MD simulations on the microsecond timescale with the CHARMM force field, and the SF adopts the crystal structure-like, inactivated conformation (PDB ID: 1K4D) [24]. Our previous work on the Kv1.2 channel had predicted aspartate flipping, SF dilation and loss of K^+^ ions from S0-S2, also using the CHARMM force field [30], before recent cryoEM structures of stably-inactivated *Shaker* W434F, Kv1.2 W362F and Kv1.3 channels became available (**Fig. 6 C-F**) [26,28,29]. In MthK, previous MD simulations done with the CHARMM force field (and reduced interaction strength between K^+^ ions and carbonyl oxygens) revealed that K^+^ ions from sites S1, S2, and S3 need to unbind before the SF collapse, which was then structurally similar to the one seen in KcsA, having a constriction at the level of the ‘first’ glycine (G61), V60 and G61 carbonyl flipping and increased number of water molecules behind the SF [31].

Here, we probed the stability of the SF of both MthK WT and the V55E in 20 independent, 5 us long unbiased MD simulations per force field/system, under two applied voltages (300 and 150 mV), resulting in 800us of sampling, and at a large activation gate opening, which speeds up SF conformational transitions [24,25]. We focused on whether the hydrogen bonds between D64 and Y51 are retained or broken (**Supplementary Fig. 2-9**, the distance larger than ca. 0.7 nm indicates a broken interaction and a subsequent aspartate flip), and on the changes in the SF geometry, at the level of both ‘first’ and ‘second’ glycines (G61 and G63, **Supplementary Fig. 10-17** and **Supplementary Fig. 18-25**, respectively).

In many of our simulations, the SF of both MthK WT and V55E undergoes dilation and/or constriction transitions at the level of G63 (and also G61), which is preceded by at least one broken D64-Y51 interaction (aspartate flip) (**Fig. 6 G-I**, **Supplementary Fig. 2-9**). Specifically, this aspartate D64 flip has been shown as a central event underlying SF inactivation in voltage-gated channels, leading to the widening of the SF at the ‘second’ glycine (**Fig. 6 C-F**). Therefore, we used G63 CA distance variations as a coordinate defining conducting and inactivated filter conformations (**Supplementary Fig. 18-25**). Based on this coordinate, we counted the conducting to non-conducting (inactivated) transitions occurring in independent simulations (**Fig. 6 J-M**), at two different voltages (150 and 300 mV). On the MD simulations timescale used here, these transitions occur irreversibly, i.e. maximally once per simulation. In the AMBER force field, we observed 7/20 such ‘inactivation transitions’ for V55E at 300 mV and 4/20 at 150 mV, whereas it did not occur at all for WT at either voltage. In the CHARMM force field, V55E transitioned to the inactivated state in 19/20 simulations at 300 mV, and 13/20 at 150 mV, whereas only 7/20 ‘inactivation transitions’ at 300 mV, and 4/20 at 150 mV, were observed for WT. These simulations clearly show that MthK V55E has an increased propensity to enter the inactivated conformation as compared to WT, in both force fields. Moreover, the inactivation rate is increased at higher voltages, in agreement with electrophysiology (**Fig. 2**). Thus, in MD simulations V55E reduces the stability of the conducting conformation of the SF and promotes inactivated conformations, which is in very good agreement with the decreased lifetimes of the open conformation seen in electrophysiological recordings (**Fig. 2**). These inactivated conformations of the SF, in both MthK WT and V55E, structurally resemble the conformations of stably-inactivated mutants of *Shaker* and Kv1.2 channels, as well as WT Kv1.3 channels (**Fig. 6 G-I**). However, they also share some similarities with filters of inactivated KcsA and MthK at a low potassium concentration, namely flipping of carbonyl groups of V60 and G61, due to water entry into the SF, as well as increased number of water molecules behind the SF (**Fig. 6 G-I, Supplementary Fig. 26**). Overall, the inactivated SFs in our simulations are quite dynamic and adopt a wide range of conformations unified however by the presence of D64 flips, widened upper parts, flipped V60 carbonyls, and a lower occupancy of the S0 ion binding site (**Supplementary Fig. 27**).

To gain more insights, we focused on molecular events happening in these MD simulations, in which D64 flipping and subsequent SF inactivation occurred, in both MthK WT and Mthk V55E (**Fig. 7**). We specifically looked at short MD trajectory fragments, just before the D64 flipping events, because, as argued before, it is an essential feature of the SF inactivation, both in our simulations and in the available structures of Kv channels. We observed that in MthK WT (for which inactivation occurred only in the CHARMM force field), there is an increased probability of water molecules entering the region behind the SF just before the aspartate flip (**Fig. 7 B**, red curve), which likely affects the stability of the Y51-D64 hydrogen bond. As explained above, this hydrogen bond is conserved in many K^+^ channels and stabilizes the D64 side chain in the non-flipped orientation (**Fig. 7 A**). At the same time, there is a slight increase in the probability of water molecules entering the SF (yellow curve) and a slight decrease in the occupancy of the top ion binding sites by K^+^ ions (purple curve).

**Figure 7.**
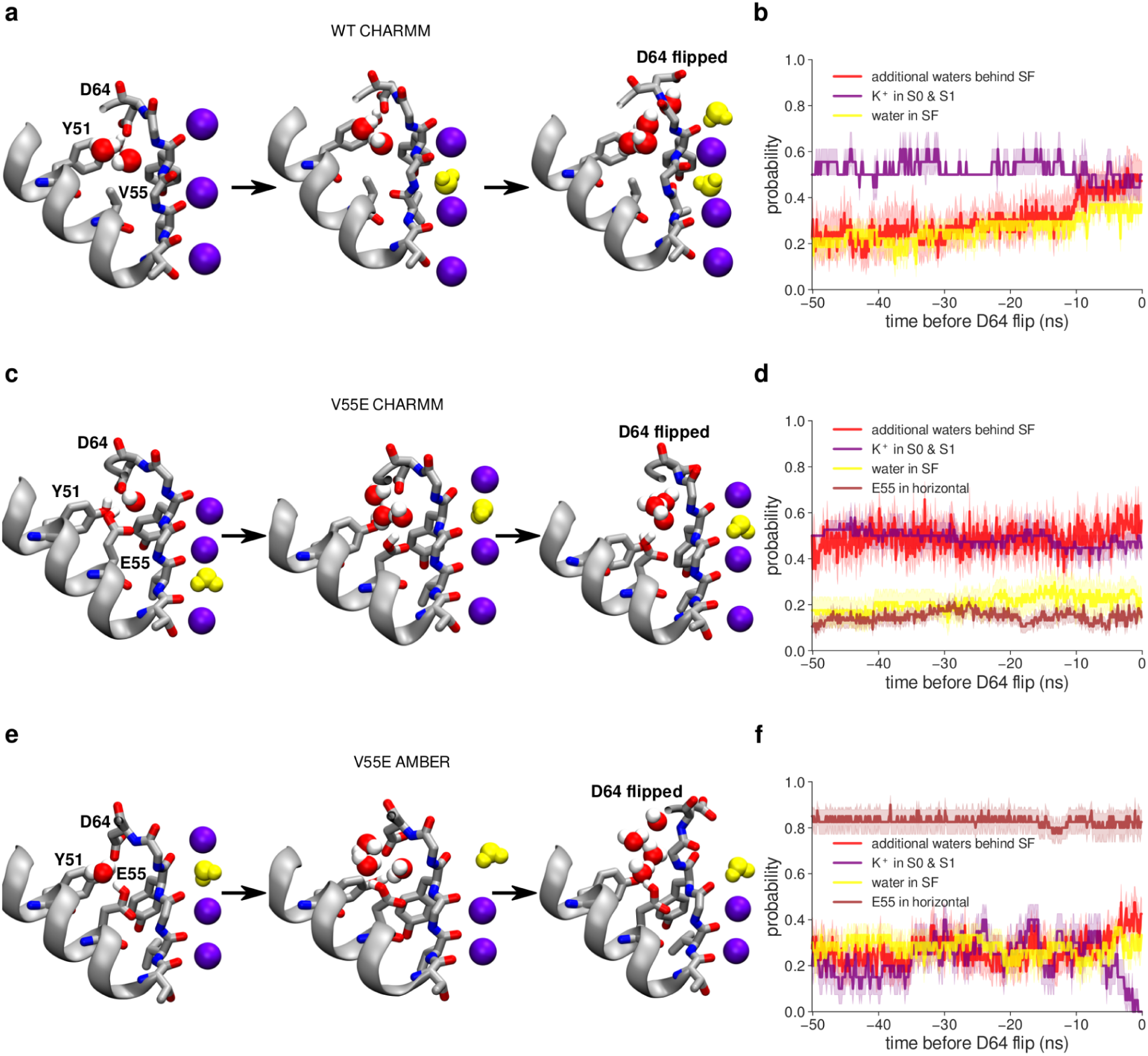
Molecular events preceding the D64 flipping and filter inactivation in MD simulations of MthK WT and V55E channels. A, C, E, Visualisations of the SF surroundings just before the D64 flip, in MthK WT simulated with the CHARMM force field, in MthK V55E simulated with the CHARMM force field, and in MthK V55E simulated with the AMBER force field, respectively. No D64 flips nor inactivations occurred in MthK WT simulated with the AMBER force field, therefore it is not included. Water molecules inside the SF are shown in yellow. B, D, F, Probabilities of specific events in 50ns simulation time before the D64 flip: additional water molecules (to the crystallographic waters) entering the space between V/E55 and D64 behind the SF (red curve), potassium ions occupying ion binding sites S0 and S1 (purple curve), water occupying inner ion binding sites (S1-S3) in the SF (yellow curve), and E55 side chain adopting the ‘horizontal’ orientation (brown curve). For calculating probabilities, all trajectories in which a D64 flip occurred were aligned at the time of the first D64 flip (t=0) and the probability was at each frame over all of these trajectories. Note that for red and brown curves, the probability is calculated for all four monomers, i.e. the value 1 would mean an event occurring in all four monomers simultaneously.,. Error bars represent the standard error.

A similar overall pattern is observed in simulations of MthK V55E with the same force field (**Fig. 7 C, D**), however the probability of water molecules entering behind the SF is higher throughout the 50ns period prior to the aspartate flip than in MthK WT (**Fig. 7 B**). We attribute this effect to the presence of a hydrophilic, protonated glutamate side chain, which has a higher propensity to attract water molecules than (hydrophobic) valine present in MthK WT. The E55 side chains adopt mostly the ‘vertical’ orientation (**Fig. 7 D**, brown curve), however the increased probability of ∼0.2 (from ∼0.1) of the E55 side chain to adopt the ‘horizontal’ orientation means that it occurs quite often in one of the monomers (on average, see the caption of **Fig. 7**), which is most likely correlated with the entry of additional water molecules behind the SF, that breaks the Y51-D64 hydrogen bond and affects the E55 side chain dynamics (**Fig. 7 C**). There are no major differences in the probability of water molecules entering the SF between MthK WT and V55E, in the CHARMM force field.

In simulations of MthK V55E with the AMBER force field (**Fig. 7 E, F**), there is a similar, high probability of water molecules entering behind the SF, and almost all four E55 side chains adopt the ‘horizontal’ orientation (brown curve) at all times. Most strikingly however, there is a sharp drop in the probability of K^+^ occupying S0 and S1 ion binding sites (purple curve), just before the D64 flipping event. This suggests that potassium ions bound to the top of the SF stabilize the conducting conformation, in the AMBER force field, with the non-flipped D64 side chains. Accordingly, control calculations in simulations of MthK WT with the AMBER force field (**Supplementary Fig. 28**), where the D64 flipping does not occur, show a consistent occupancy of the S0 and S1 by potassium ions.

Summarizing, we discovered that the V55E mutation reduces the stability of the conducting conformation of the SF in the MthK channel, by promoting water entry behind the SF, leading to the breakage of the Y51-D64 hydrogen bond and subsequent flipping of the D64 side chains. This entry of additional water molecules likely correlates with the ‘horizontal’ orientation of the E55 side chain. Even though both force fields predicted an overall similar effect of the V55E on the SF stability, there are important differences. Firstly, as already described for shorter simulations, the AMBER force field strongly prefers the ‘horizontal’ orientation of the E55 side chain, due to the differences in dihedral parametrization (**Supplementary Fig. 1**). Secondly, the distribution of K^+^ ions in the SF, and the effect of the mutation on the overall number of K^+^ ions in the SF, vary between the force fields (**Supplementary Fig. 27**). Importantly, the D64 flipping in MthK V55E simulated with the AMBER force field strongly correlates with the unbinding of K^+^ ions from the top of the SF. Thirdly, the same aspartate flipping and inactivation mechanism as in MthK V55E is seen in simulations of MthK WT, but only when the CHARMM force field is used.

We further investigated the possible reasons for the last discrepancy. We evaluated two factors, possibly affecting the D64 side chain dynamics: i) the strength of the Y51-D64 hydrogen bond (weaker interactions would promote the aspartate flip, **Supplementary Fig. 29 A-C**); and ii) the energetics of dihedrals defined by the heavy atoms of the D64 side chain (high energy orientations of the D64 side chains would contribute to a lower stability of a given state, **Supplementary Fig. 29 D-F**). From this analysis, the differences between the two force fields become more clear - surprisingly, the Y51-D64 interactions are actually stronger in CHARMM than in AMBER (opposed to the inactivation rate trend). Strikingly however, one of the two dihedrals of the D64 side chain has a high energy (up to ca. 19 kJ/mol) in the CHARMM force field, when the D64 side chain is in the non-flipped orientation, and is able to adopt a lower energy conformation only after the aspartate flip (**Supplementary Fig. 29 E**). In stark contrast, in AMBER simulations, both dihedrals are low in energy when the D64 side chain is in the non-flipped orientation (**Supplementary Fig. 29 F**). These force field differences, primarily in the dihedral parametrization, resulting in different preferred orientations of the D64 side chain, and, to some degree, in the K^+^ occupancy of the individual ion binding sites (high S0 occupancy in AMBER, **Supplementary Fig. 27**), likely explain the much higher propensity of the MthK WT SF inactivation in MD simulations with the CHARMM force field. Interestingly, these inactivated conformations resemble experimentally solved structures of Kv channels (**Fig. 6**).

We wondered how these two factors are affected by the V55E mutation. The energetics of the dihedrals is only slightly affected, in both force fields (**Supplementary Fig. 30**), although the high energy dihedral in CHARMM is even higher in energy in MthK V55E. The energetics of the Y51-D64 and E55 (or water in MthK WT)-D64 hydrogen bonds is also affected, especially in the CHARMM force field: most of these interactions are stronger in MthK WT than in the V55E mutant (**Supplementary Fig. 30 C**), which could also contribute to the lower stability of the conducting state in MthK V55E in this force field. In the AMBER force field, the trend is not so clear: some interactions are stronger in MthK WT and some in MthK V55E. Therefore, in MD simulations of MthK V55E with the AMBER force field, we assign the unbinding of K^+^ ions from the top of the SF, and the entry of additional water molecules behind the SF as the primary cause of the aspartate flipping and SF inactivation.

### Free energy calculations - E55 pKa

In our MD simulations we have repeatedly observed (mostly in the AMBER force field) transitions of the E55 side chain from the ’vertical’ orientation, where it closely interacts with D64, to the ‘horizontal’ orientation, without a clear interaction partner, and instead interacting with a water molecule (or causing an entry of a larger number of water molecules). The question arises whether the E55 side chain would still stay protonated in the horizontal orientation. This putative deprotonation of the E55 side chain would presumably have a substantial effect on all structural observables in MD simulations. Therefore, to assess the likelihood of the E55 side chain to deprotonate, we used non-equilibrium free energy calculations to calculate the *ΔG* of E55 deprotonation and consequently its *pKa.* (**Supplementary Fig. 31**, see Methods) [32–35]. Free energy results in both force fields are in good agreement, showing the *ΔG* of E55 deprotonation of more than 40 kJ/mol (with respect to the reference state in bulk), indicating that it is unlikely for E55 to be in the deprotonated form. In fact, such a value translates to a shift of the *pKa* (*ΔpKa*) of 7-8 units from the glutamate experimental value of 4 [36]. Even taking into account possible force field and simulation inaccuracies, these results strongly suggest that E55 stays protonated at all times.

## DISCUSSION

In this work, we set out to better understand the molecular basis for SF gating in MthK WT and the KcsA-like mutant MthK V55E by combining electrophysiological measurements with large scale MD simulations. Previously, it was established that MthK WT exhibits SF gating at depolarized voltages, similar to C-type inactivation described in KcsA and Kv channels [9,12]. Here, both electrophysiology and computational methods showed that in MthK V55E, the open state of the channel is destabilized. Gating of V55E in experiments is characterized by markedly reduced Popen over a wide range of transmembrane voltage (−200 to 200 mV), and at high potassium concentration (200 mM) at both sides of the membrane, as opposed to MthK WT, in which the open state is destabilized depolarized voltages and/or reduced external potassium concentration [9,12]. Moreover, V55E shows markedly lower single channel unitary conductance in electrophysiological experiments than MthK WT.

Two *in silico* electrophysiology simulation-based approaches revealed how the V55E mutation affects ion permeation and the SF stability in MthK on the atomistic scale. In contrast to the KcsA crystal structure, where the side chain of this protonated glutamate is thought to exist only in one rotameric state (‘vertical orientation’), hydrogen bonding with a conserved aspartate and crystallographically resolved water molecule behind the SF, our MD simulations suggest that in MthK V55E, the E55 side chain is mobile and can exist in at least two rotameric states, namely ‘vertical’ and ‘horizontal’. The strong interaction between E55 and D64 in the ‘vertical’ orientation leads to decreased structural plasticity of the SF in its upper part, which increases energetic barriers for ion permeation (in the AMBER force field) and compromises the geometry of the S0 binding site (in the CHARMM force field), resulting in much lower simulated outward currents through MthK V55E. This strain in the SF is relieved when the E55 side chain adopts the ‘horizontal’ orientation, and then the mutant channel is able to permeate K^+^ ions at rates approaching those recorded in MthK WT. Therefore, the ionic current in MthK V55E is regulated not only *via* activation gate-SF coupling (as previously described for MthK WT) [21], but also by the E55 side chain rotameric state. The preference for the ‘vertical’ vs ‘horizontal’ orientation also depends on the level of the activation gate opening, revealing a complex activation gate-selectivity filter-pore helix coupling. Further, the two molecular force fields used in this work also predicted a very different propensity of the E55 side chain to adopt the ‘horizontal’ orientation: in the AMBER force field, it was actually the preferred orientation in many cases, whereas it in the CHARMM force field it was hardly accessible, especially at the shorter timescales.

Our longer simulations also allowed us to peek into larger conformational transitions of the SF, which we relate to the C-type inactivation process. Here, surprisingly, both MthK WT and MthK V55E show behavior more comparable with the stably-inactivated Kv channels (*Shaker* W434F, Kv1.2 W362F and WT Kv1.3), although some features seen with in KcsA, such as carbonyl flipping of V60 and G61 residues, as reported before [31], are present as well. These observations suggest that the differences between MthK and KcsA channels are not only limited to the residue at the position 55 in MthK (valine in MthK WT, protonated glutamate in KcsA), and likely the variation in other residues in the pore-helix play a major role as well in regulating the SF gating behavior of these channels. Indeed, the distribution of hydrophobic and hydrophilic residues vary in the pore region of both channels (**Supplementary Fig. 32**). One major difference is the interaction partner of the aspartate behind the SF: in MthK it is a tyrosine residue (Y51), whereas in KcsA it is a tryptophan residue (W67). These residues have different hydrophobicity and hydrogen bonding capabilities, therefore they might largely affect the behavior of the aspartate side chain, and subsequently its other interaction partner, namely protonated glutamate (in KcsA and in MthK V55E). The double MthK Y51W,V55E mutant would be an interesting construct in the future work, to see if it shows a behavior even more related to KcsA. Another interesting difference is position 64 in KcsA (positively charged arginine), which in MthK is replaced by valine (V48). A positively charged residue near D80 might stabilize the aspartate in the non-flipped orientation. Residues that form the extracellular loop following the SF have also been postulated to play a role in C-type inactivation in KcsA. For example, the entry of water molecules behind the SF in KcsA was proposed to occur through the Y82 ‘lid’ residue [37,38]. MthK has a serine residue in this position, and tyrosine occurs one position earlier in the sequence. These variations in the extracellular loop might underlie different water penetration pathways in KcsA and MthK, again leading to the divergent dynamics of D80 and other residues behind the SF.

Similar asymmetric SF conformations to ones reported in this work have been also observed in the structures of some two-pore domain (K2P) potassium channels: TREK-1 [39] and TASK-2 [40]. Specifically, we highlight that the conformational transitions (inactivation) observed in our MD simulations of both MthK WT and V55E channels involve widening (dilation) of the SF at the level of ‘second’ glycines (G63 in MthK), in agreement with available structural data for *Shaker* W434F, Kv1.2 W362F and WT Kv1.3 channels [26,28,29].

Our simulation data reveal that this dilation follows the breakage of the critical hydrogen bond behind the SF, between D64 and Y51. The strength of this hydrogen bond has been previously directly linked to the rate of inactivation in Kv channels in experiments employing non-natural amino acids [10], further highlighting the converging behavior of MthK and Kv channels with respect to SF gating. When this Y51-D64 hydrogen bond is broken (e.g. via the entry of additional water molecules behind the SF), the D64 side chain flips toward the extracellular side (aspartate flip) allowing the SF to widen. Interestingly, a combined experimental and computational study on the ‘MthK-like’ KcsA E71V mutant showed a distinct mode of SF inactivation, without the filter constriction typical for KcsA WT, nor filter dilation, but rather with the overall rigidified filter [41]. We note however that in the structures of KcsA E71V at large gate openings, the hydrogen bond between the aspartate and tryptophan (analogous to the Y51-D64 bond in MthK) was broken [41], although a ‘full’ aspartate flip, with its side chain pointing toward the extracellular side, was not reported. On the other hand, the structures of (non-inactivating) KcsA E71A at low potassium concentration show aspartate and valine flipping in the SF, akin to features reported in this work [42].

In our simulations, the widened SF displays compromised geometries of the ion binding sites, due to outward rotation of carbonyl oxygens as reported earlier [12,42,43]. Subsequently, the ion binding sites get filled with water molecules (**Fig 6, G-I, Supplementary Fig. 26**), which disrupts the optimal ion permeation that normally occurs in a water-free fashion *via* direct ion-ion contacts [44]. By directly simulating many inactivation events (160 individual simulations in total), we showed that this process occurs more frequently in MthK V55E than in the WT channel and is accelerated by more positive voltages, in full agreement with our electrophysiological recordings. Interestingly, our data suggest that the structural basis for reduced stability of the MthK V55E SF is, again, related to the orientation of the E55 side chain. As E55 participates in the intricate hydrogen bond network including D64 and Y55, the rotameric transition of its side chain from ‘vertical’ to ‘horizontal’ leads to the entry of additional water molecules behind the SF, further destabilizing D64-Y55 bond. In the ‘horizontal’ orientation, the hydrophilic side chain of protonated E55 does not have any obvious hydrogen bond partners among protein residues to bond with, and it attracts additional water molecules from the extracellular side. Viral K^+^ channels, which also exhibit exquisite SF gating, have hydrophilic threonine or serine residues in the position equivalent to E55 [45,46], suggesting that a similar molecular behavior might be expected in those channels.

We note major differences between the used force fields: CHARMM and AMBER, in their preferences regarding the ‘vertical’ and ‘horizontal’ orientations of the E55 side chain, as well in the flipped and non-flipped orientations of the D64 side chain. We uncovered the origins of these differences being different parameterizations of dihedrals of these residues, and further work is needed to resolve this issue. It is interesting that the ‘horizontal’ orientation of the E55 side chain (or E71 in KcsA) has not been seen before, neither in structural work nor in MD simulations. We suggest the following reasons for this situation. First, KcsA channels have been mostly simulated with the CHARMM force field (with some recent exceptions [47]), in which the side chain of protonated E71 has a much lower propensity of adopting the ‘horizontal’ orientation. Second, our analysis suggests that the ‘horizontal’ orientation is favored for the larger openings of the activation gate. Most of the KcsA structures have been solved in the closed state, and in those solved in the open state, protonated glutamate is replaced by alanine (the E71A construct), to remove C-type inactivation. Third, as discussed above, MthK V55E and KcsA channels are quite different in their pore region (**Supplementary Fig 32**), thus it is possible that the ‘horizontal’ orientation is more common for MthK V55E than for KcsA. Our work is the first dealing with the MthK V55E channel, for which there is currently no structural data. We cannot however fully rule out a possibility that the high frequency of the ‘horizontal’ orientation is an artifact of the AMBER force field, and we recommend further work in this direction.

In our simulations, the inactivation process in the MthK channel, arising from SF gating, seems to be structurally related to the same process in Kv channels, but with some characteristic shared with KcsA. By combining electrophysiology and MD simulations, we showed that a single mutation in the pore-helix (V55E) can simultaneously affect conducting properties and open state stability, enhancing the intrinsic propensity of WT channels to inactivate. We propose a mechanism of this effect based on a functionally relevant rotameric transition of a hydrophilic residue behind the SF, coupled with a water entry behind the SF and potassium ion unbinding from the top of the filter. These results contribute to a conceptual framework for the effects of pore-helix residues on SF gating in various potassium channels.

## METHODS

### Protein preparation and single-channel recording

MthK was expressed, purified, and reconstituted into proteoliposomes as described previously [48]. Proteoliposomes were composed of *E.coli* lipids (Avanti) and rapidly frozen in liquid N_2_ and stored at −80°C until use. Protein concentrations in proteoliposomes ranged from 5 to 25 μg protein per mg lipid. Mutations were introduced using the QuikChange Site-Directed Mutagenesis Kit (Agilent Technologies) and confirmed by DNA sequencing.

Recordings were obtained using planar lipid bilayers of POPE:POPG (3:1) in a horizontal bilayer chamber, essentially as described previously [12]. For these experiments, solution in the *cis* (top) chamber contained 200 mM KCl, 10 mM HEPES (pH 8.1), 100 µM Cd^2+^. Solution in the *trans* (bottom) chamber contained either 5 or 200 mM KCl and 10 mM HEPES (pH 7.0). This arrangement ensured that MthK channels incorporated into the bilayer with the cytosolic side facing the *trans* chamber would be silent, whereas channels with the cytosolic side facing the *cis* chamber would be maximally activated (by Cd^2+^). Within each bilayer, multiple solution changes were performed using a gravity-fed perfusion system that enabled exchange of solution in the *trans* chamber. To ensure completeness of solution changes, the *trans* chamber was washed with a minimum of 10 ml (∼10 chamber-volumes) of solution prior to recording under a given set of conditions.

Single-channel currents were amplified using a Dagan PC-ONE patch clamp amplifier with low-pass filtering to give a final effective filtering of 1 kHz. This level of filtering corresponds to a dead time (minimum time required for an event to reach 50% threshold) of 0.179 ms. Currents were digitized at a rate of 50 kHz (20 µsec per sample) and analyzed by measuring durations of channel openings and closings at each current level by 50% threshold analysis, using pClamp 9.2. These were used to calculate NPo as:

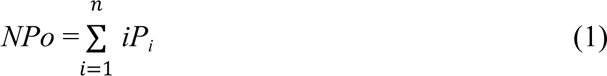

in which *N* is the number of channels in the bilayer, *i* is the open level and *P_i_* is probability of opening at that level. The mean single-channel open probability (*Po*) is obtained by dividing *NPo* by *N*, determined by recording under conditions where the maximum level of channel opening can be observed. The voltage-dependence of *Po* was described by the Boltzmann equation,

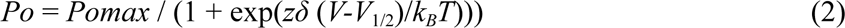

in which *Pomax* is the maximal *Po*, *zδ* is the effective gating valence (in e_0_), *V_1/2_* is the voltage at half-maximal *Po*, *k_B_* is Boltzmann’s constant and *T* is temperature. Data points (e.g. *Po* and mean interval durations) are presented as mean ± SEM of three to five observations for each data point, and collectively represent data from a total of 17 different bilayers.

Mean unitary current amplitudes were determined by constructing all-points histograms from digitized currents at each voltage. These could be described in part by a sum of two Gaussian distributions,

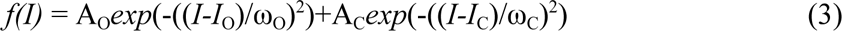

in which A_O_ and A_C_ are the heights of the distributions of open and closed current levels, *I*_O_ and *I*_C_ are the means of the distributions, and ω_O_ and ω_C_ are the widths of the distributions. Thus the unitary current amplitude is given by the difference *I*_O_ minus *I*_C_. As a check on this method of estimating mean current amplitudes, amplitudes were additionally determined for individual idealized openings in conjunction with 50% threshold analysis using pClamp9.2. In this approach, the open amplitude is calculated as the difference between the closed baseline level and mean current level following a 50% threshold crossing, excluding points during the transitions between the open and closed levels (*i.e.* 0.179 ms after the threshold-crossing for opening, and 0.179 ms prior to the threshold-crossing for closing for currents in this paper, which were low-pass filtered at 1 kHz, as described in the pClamp User Guide). If the measured duration of the opening is less than 0.358 ms (i.e. 2 x 0.179 ms), then the amplitude is recorded as the current level at the midpoint of the opening.

### MD simulations

We used the same setup as we used for MthK WT in our recent publication [21], where we thoroughly characterized ion permeation and selectivity filter-activation gate coupling, using the CHARMM36m (referred to as CHARMM) [49,50] and AMBER14sb (referred to as AMBER) [51] force fields. We did not use any further modifications of the carbonyl oxygen - potassium ion interactions, as non-modified force fields seem to better reproduce solvation free energies [52].

The data for MthK WT in the current manuscript (black traces) are taken directly from our previous work [21]. For MD simulations of the V55E mutant, the valine residue in MthK WT was replaced by protonated glutamate from KcsA (PDB ID: 1K4C) after aligning both channels with their (identical) selectivity filters. The resulting structure was further equilibrated for at least 100 ns, whereas all other simulation details and parameters were identical to those used before. After equilibration, we imposed the desired degree of activation gate opening by applying harmonic distance restraints between the CA atoms of the pairs of terminal residues - P19 and F97. The equilibrium distances were adopted from our previous publication, and the resulting average distances are plotted in **Fig. 4 B** and **Fig. 5 A**. For additional simulation sets, where the orientation of the E55 side chain was restrained as well, additional restraints were used. In the AMBER force field, where the ‘horizontal’ orientation is frequently visited, we used the following restrains: to keep the E55 side chain in the ‘vertical’ orientation, we restrained the distance between the proton from the protonated side chain of E55 and one of the carboxylate oxygens of the D64 residue, to the distance of 0.167 nm (an average seen in non-restrained simulations), with a force constant of 500 kJ/mol/nm^2^. To keep the E55 side chain in the ‘horizontal’ orientation, we restrained the distance between the non-protonated oxygen from the E55 side chain and the amide proton of the G61 residue, that is nearby, to the distance of 0.200 nm with the force constant of 500 kJ/mol/nm^2^. In the CHARMM force field, where the ‘vertical’ orientation is seen almost exclusively, we had to use much stronger restraints to generate MD simulations with the ‘horizontal’ orientation. We first used some of the snapshots from the AMBER simulations, with the E55 side chain in the ‘horizontal’ orientation, converted them to the CHARMM force field, and then used position restraints on the side chain atoms, with a force constant of 20000 kJ/mol/nm^2^. Only such a strong force constant prevented the side chains from returning to the vertical orientation.

The systems were simulated at an external field applied along the z-axis to generate the membrane voltage of ∼300 mV. The voltage (*V*) was calculated with:

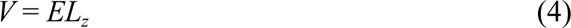

Where *E* denotes the applied electric field and *L_z_* the length of the simulation box along the z-axis. All simulations were performed with GROMACS MD software, versions 5.1 and 2020 [53–57]. All the simulation settings were identical to our previous work [21]. For each opening and force field, 10 (short simulations) or 20 (long simulations) were performed, each lasting 1 us (short simulations) or 5 us (long simulations), which resulted in a total simulation time (for all systems) of ca. 920 us. Ionic currents were estimated by counting the number of K^+^ ions crossing the SF in each simulation, using a custom FORTRAN code (available as a Supplementary Software associated with our previous publication [21]). Remaining quantities were calculated using GROMACS tools - gmx distance (distances), gmx angle (angles) and gmx select (number of water molecules). Most of the presented data (besides individual simulation traces and counts) are averages from at least 10 independent simulation replicas. Error bars represent 95% confidence intervals or SEM. For data analysis, Python 3 was used, together with numpy [58], pandas [59], seaborn, matplotlib [60], scipy [61] and bootstrapped modules. Molecular visualizations were rendered with VMD [62].

### Protonation free energy calculations

To assess the free energy of deprotonation of the E55 side chain in its horizontal orientation, we used non-equilibrium free energy calculations using GROMACS 2020 and the pmx package [33] to calculate the *pKa* shift (*ΔpKa*) of the E55 side chain from the experimental value of 4 of glutamate in bulk water [36], following a recent application [35]. In detail, one of the four protonated E55 was selected, and the proton on the carboxylate group was alchemically switched to a dummy atom. At the same time, an opposite process, i.e. switching a dummy atom to a proton on the carboxylate group was performed for glutamate-containing tripeptide (G-E-G) in bulk, that served as a reference state see **Supplementary Fig. 31 A**). Both processes were carried out simultaneously in the same simulation box and coupled to the same lambda variable, ensuring the electrical neutrality of the simulation box at all times. This way, the calculated ΔG (**Supplementary Fig. 31 D, E**) is equal to *ΔG_protein* - *ΔG_referenc*e, with the G-E-G tripeptide as a reference state, and can be directly plugged into the equation [32]:

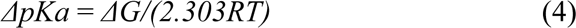

Where *R* and *T* are the gas constant and simulation temperature, respectively. In the free energy calculations, a lower temperature of 298K and salt concentration of 150 mM was used. The side chain of E55 was kept restrained with position restraints in the horizontal orientation as described above. The central atom of the G-E-G peptide was position restrained as well, to keep the tripeptide away from the membrane and the channel. For each force field, five independent equilibrations, each 20ns long for both end states, was performed. From each equilibration, a hundred equally spaced snapshots were selected (ignoring the first 5ns) for fast, non-equilibrium transitions (500 transitions from state 0 (dummy) to 1 (proton) and 500 transitions from state 1 to 0). Each non-equilibrium transition lasted 10ns, due to the relatively slow dynamics of the D64 side chain, which could flip toward the extracellular space when E55 was deprotonated (**Supplementary Fig. 31 B, C**). The final free energy difference was then calculated using pmx. Low error bars (**Supplementary Fig. 31 D, E**) across the five repeats indicate converged simulations.

Supplementary Material consists of Supplementary Figures 1-32.

## ACKNOWLEDGEMENTS

We thank Simon Bernèche and Florian Heer for many important insights and all three anonymous referees, whose comments greatly improved the manuscript, Vytas Gapsys for his help with the free energy calculations and Lucie Delemotte for the critical reading of the revised manuscript. This research was supported by the German Research Foundation (DFG) through FOR2518 ‘Dynion’, Project P5 (W.K. and B. L. dG.) and National Institutes of Health grant R01 GM126581 (B.S.R.).

## AUTHOR CONTRIBUTIONS

B.S.R., W.K. and B. L. dG. conceived the project. A.S.T. and B.S.R. performed and analyzed the electrophysiological recordings. W.K. performed and analyzed MD simulations. W.K. B.S.R. and B. L. dG. wrote the manuscript.

## Notes

### Competing Interest Statement

The authors have declared no competing interest.

### Summary of Updates

Completely new Figure 3 with more detailed analysis of single current traces. Additional data points in the Figure 4.

