## Supplementary Figures for "Driving Forces underlying Selectivity Filter Gating in the MthK Potassium Channel"

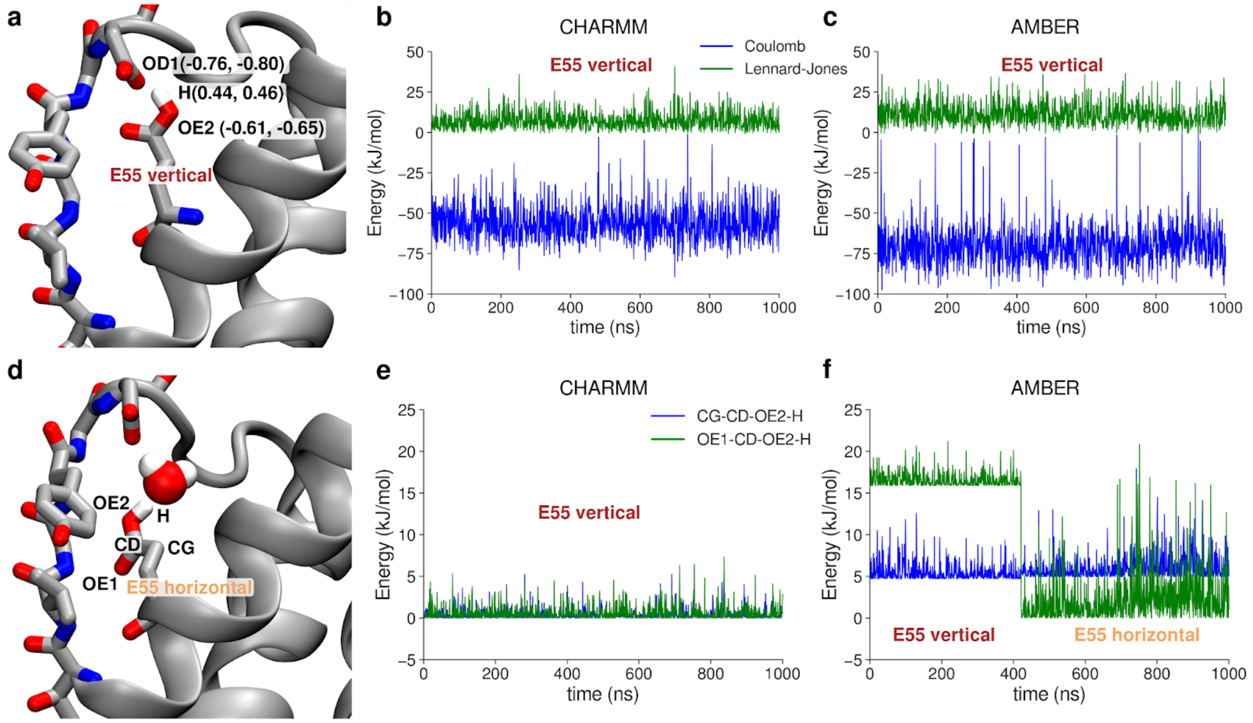


**Supplementary Figure 1 Differences in energetics of the 'vertical' vs. 'horizontal' orientation of the E55 side chain in CHARMM and AMBER force fields.** A, E55 side chain in the 'vertical' orientation, making a hydrogen bond with D64. Partial charges of atoms participating in the hydrogen bond are given in parentheses, first the value for CHARMM and second for AMBER. B-C, Exemplary interaction energies between hydrogen bond donors (H & OE2 atoms) and acceptor (OD1 atom), in both CHARMM and AMBER, respectively. D, E55 side chain in one of the 'horizontal' orientations visited in the AMBER force field. Atoms participating in two dihedral angles involving the H atom from the protonated E55 side chain are listed. E, F, Exemplary torsional energy in a trajectory of two dihedrals involving the H from the E55 side chain, in both CHARMM and AMBER, respectively.

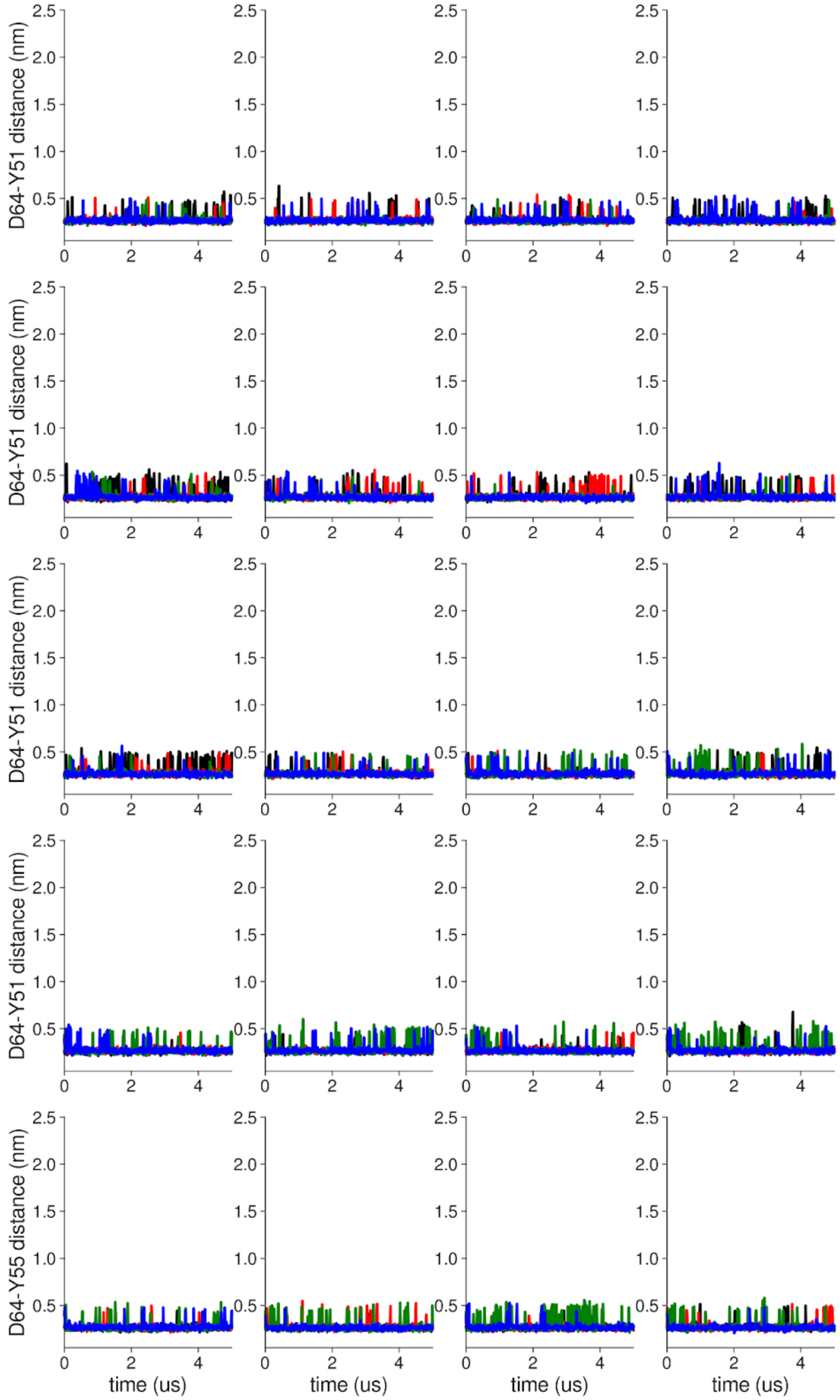


**Supplementary Figure 2 Individual distance traces between D64 CG atom and Y51 HH atom in long MthK WT AMBER simulations at 300 mV.** Each color refers to one of the channel monomers. Each panel shows traces from an independent, 5us long simulation. Distance values > 1 nm indicate broken interactions.

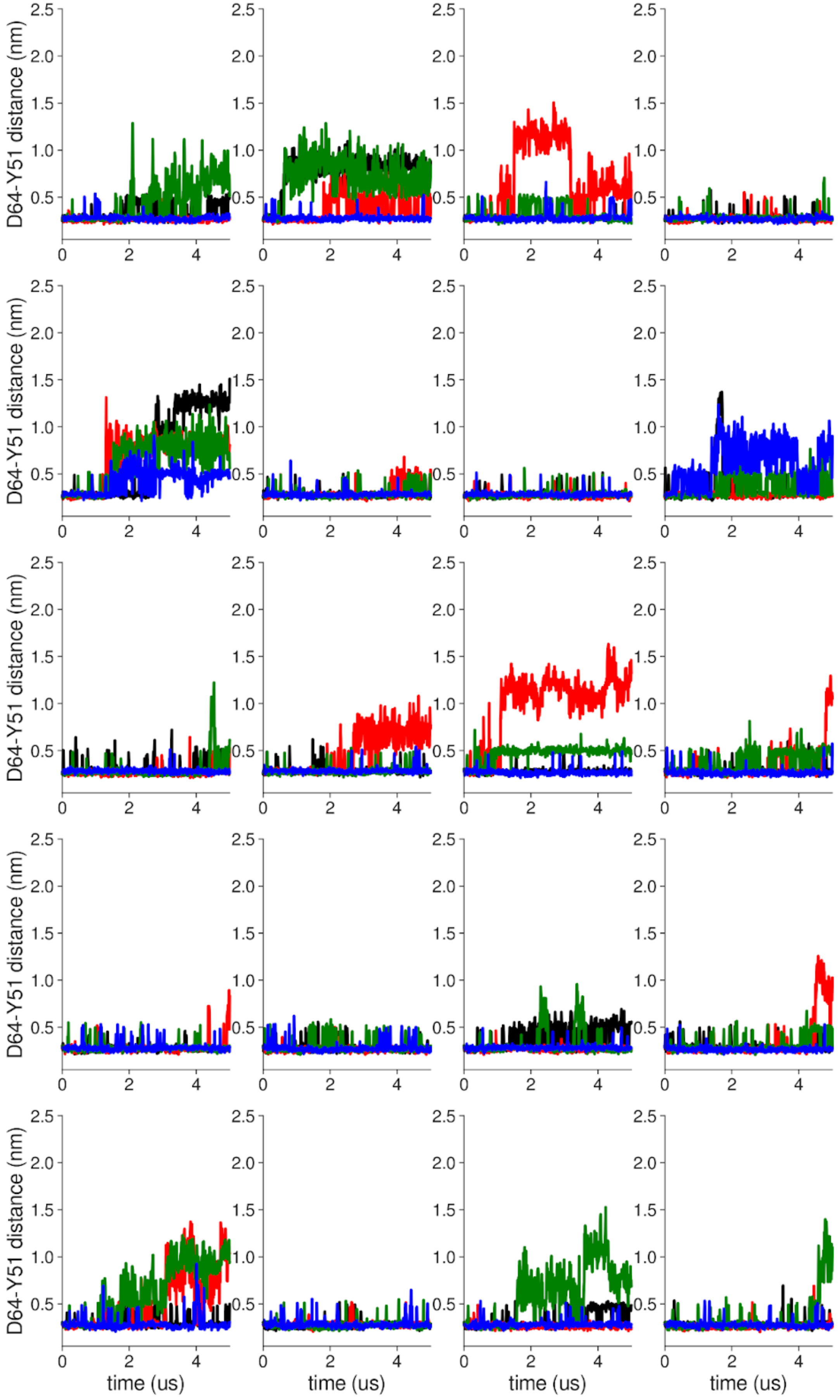


**Supplementary Figure 3 Individual distance traces between D64 CG atom and Y51 HH atom in long MthK V55E AMBER simulations at 300 mV.** Each color refers to one of the channel monomers. Each panel shows traces from an independent, 5us long simulation. Distance values > 1 nm indicate broken interactions.

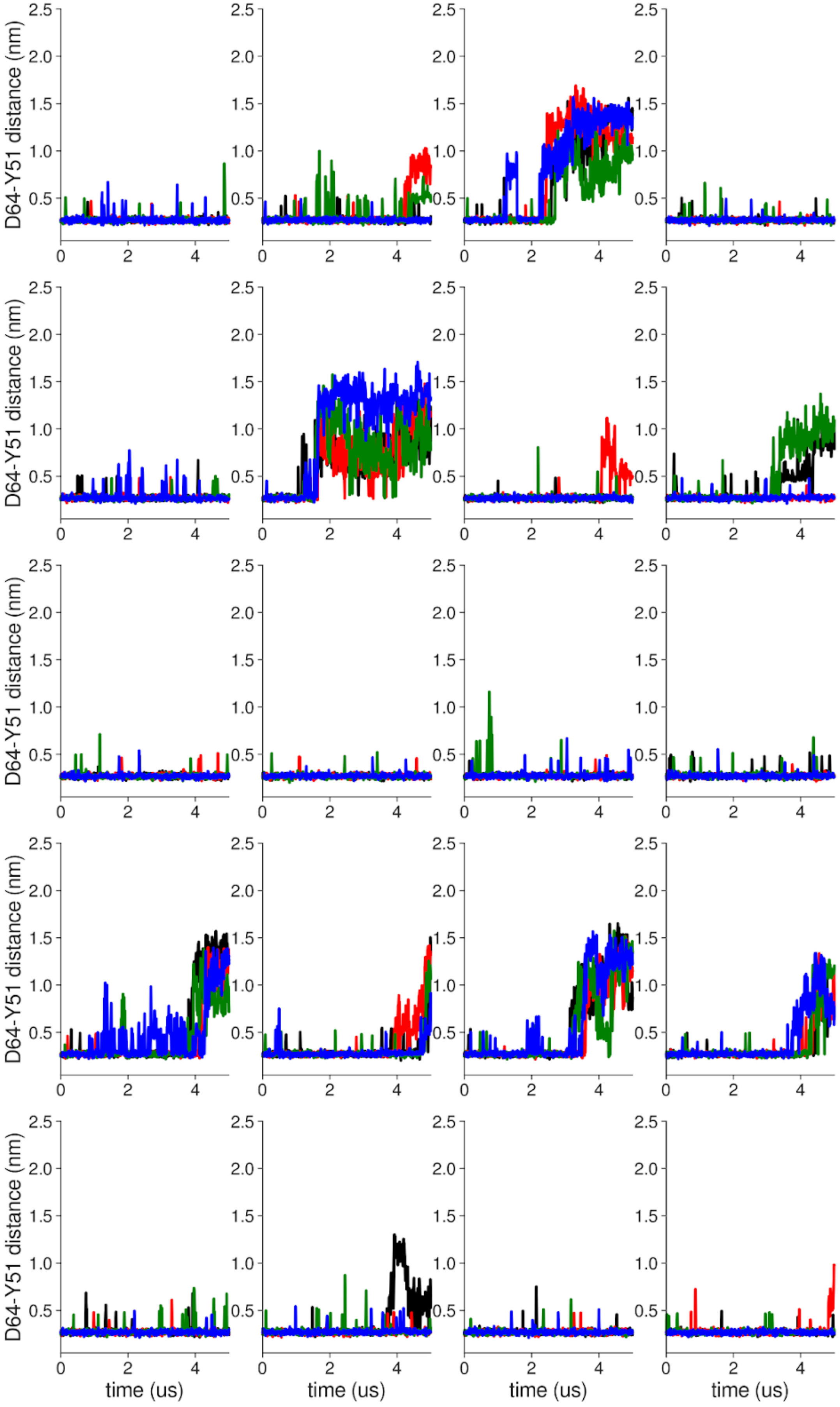


**Supplementary Figure 4 Individual distance traces between D64 CG atom and Y51 HH atom in long MthK WT CHARMM simulations at 300mV**. Each color refers to one of the channel monomers. Each panel shows traces from an independent, 5us long simulation. Distance values > 1 nm indicate broken interactions.

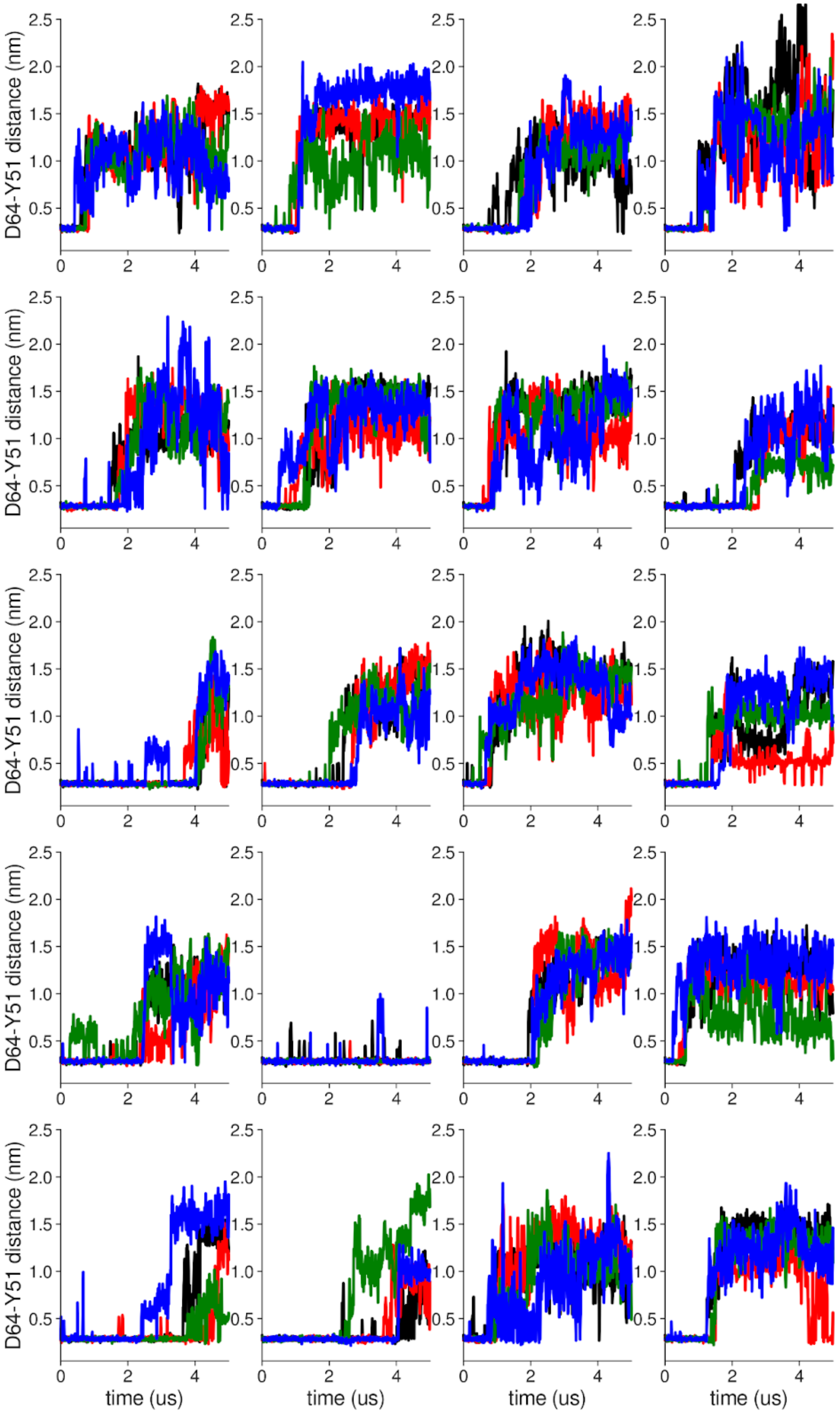


**Supplementary Figure 5 Individual distance traces between D64 CG atom and Y51 HH atom in long MthK V55E CHARMM simulations at 300 mV.** Each color refers to one of the channel monomers. Each panel shows traces from an independent, 5us long simulation. Distance values > 1 nm indicate broken interactions.

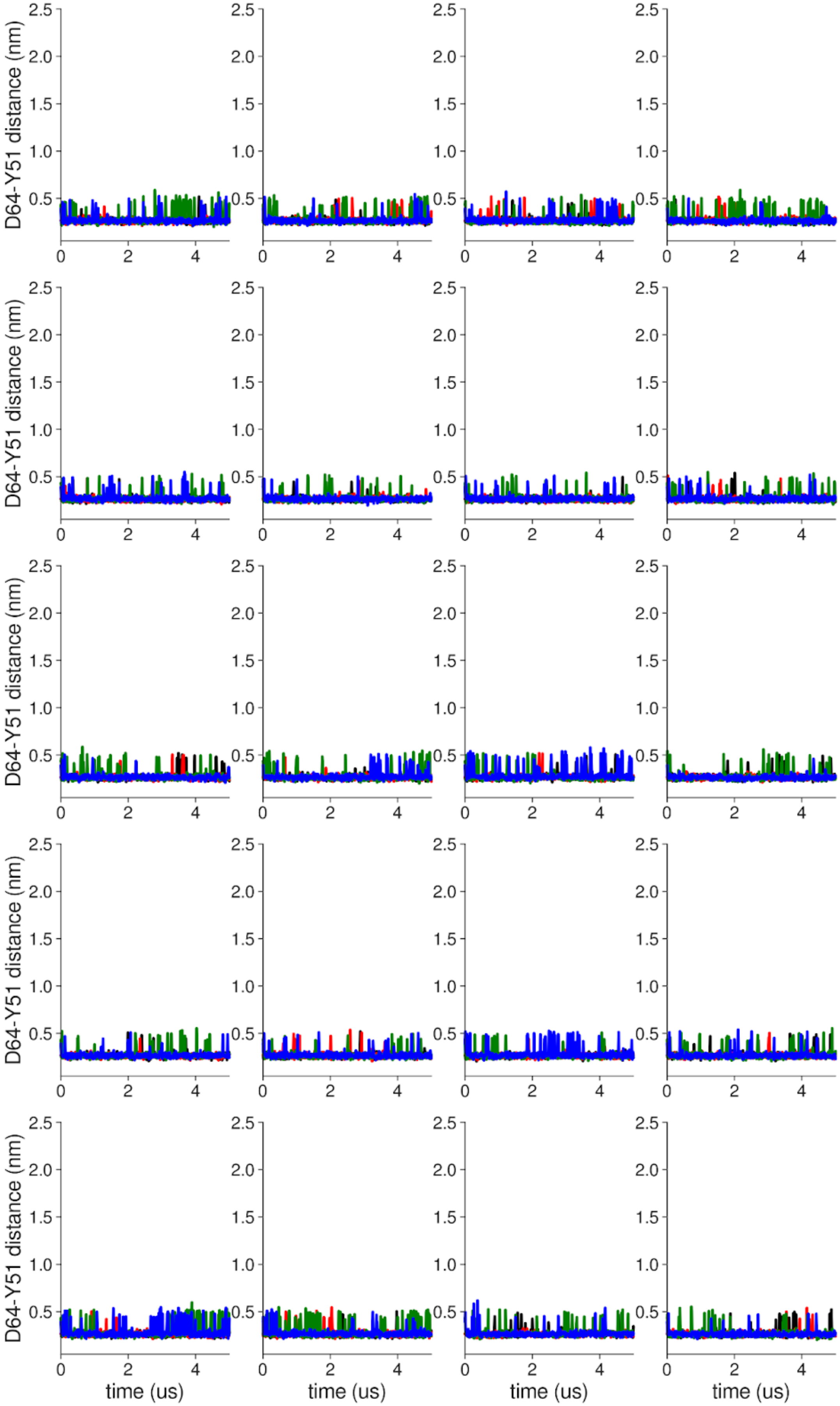


**Supplementary Figure 6 Individual distance traces between D64 CG atom and Y51 HH atom in long MthK WT AMBER simulations at 150 mV.** Each color refers to one of the channel monomers. Each panel shows traces from an independent, 5us long simulation. Distance values > 1 nm indicate broken interactions.

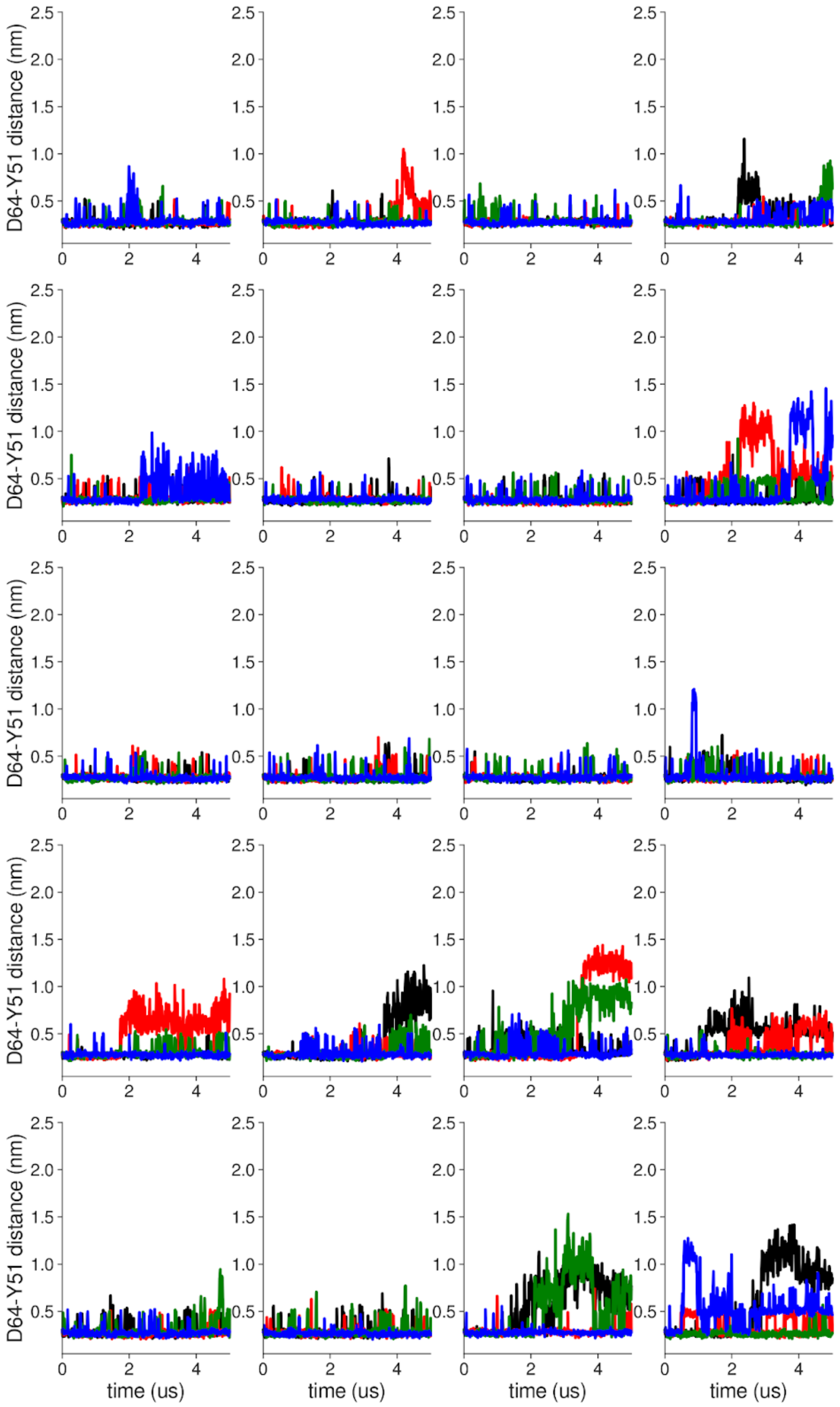


**Supplementary Figure 7 Individual distance traces between D64 CG atom and Y51 HH atom in long MthK V55E AMBER simulations at 150 mV.** Each color refers to one of the channel monomers. Each panel shows traces from an independent, 5us long simulation. Distance values > 1 nm indicate broken interactions.

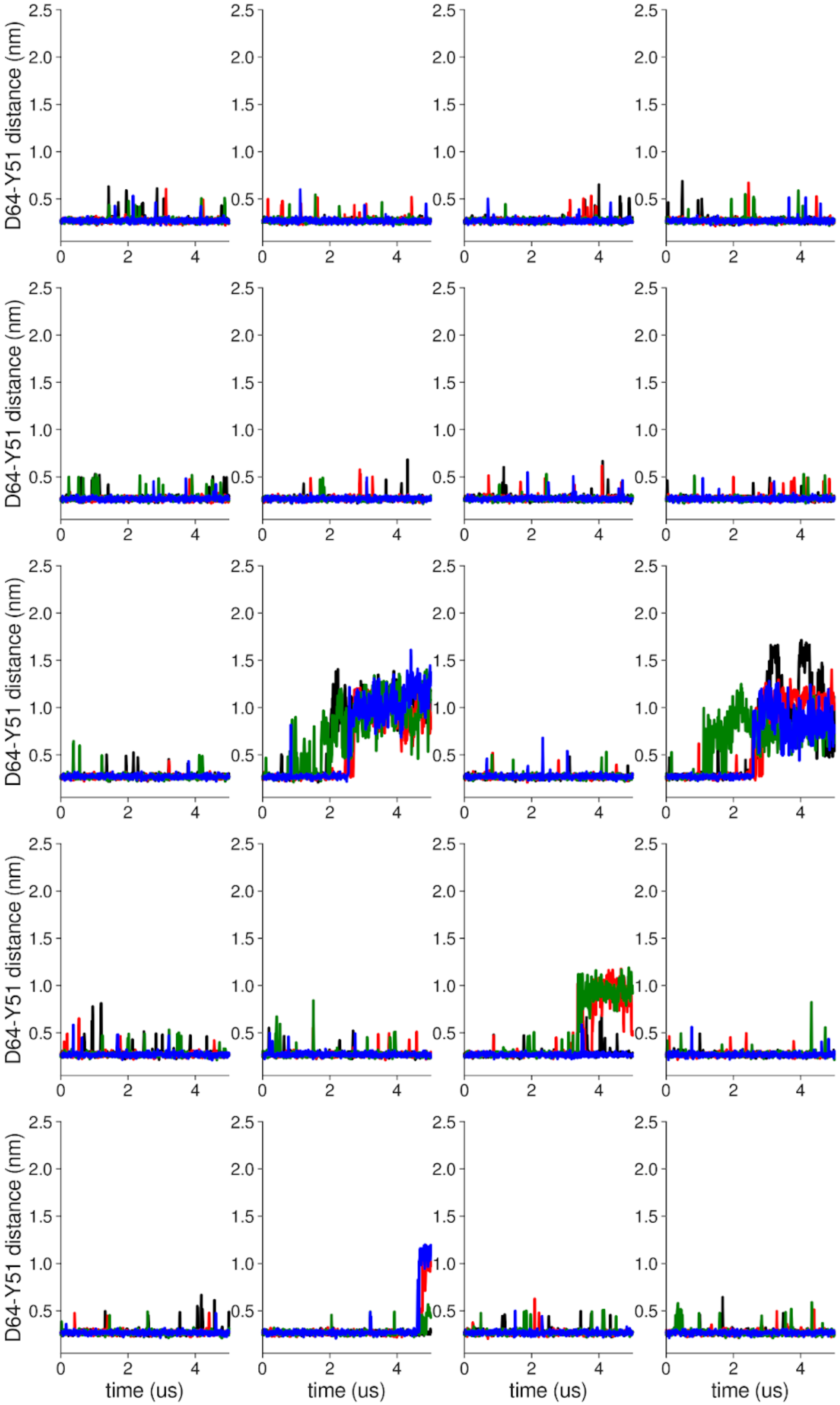


**Supplementary Figure 8 Individual distance traces between D64 CG atom and Y51 HH atom in long MthK WT CHARMM simulations at 150 mV**. Each color refers to one of the channel monomers. Each panel shows traces from an independent, 5us long simulation. Distance values > 1 nm indicate broken interactions.

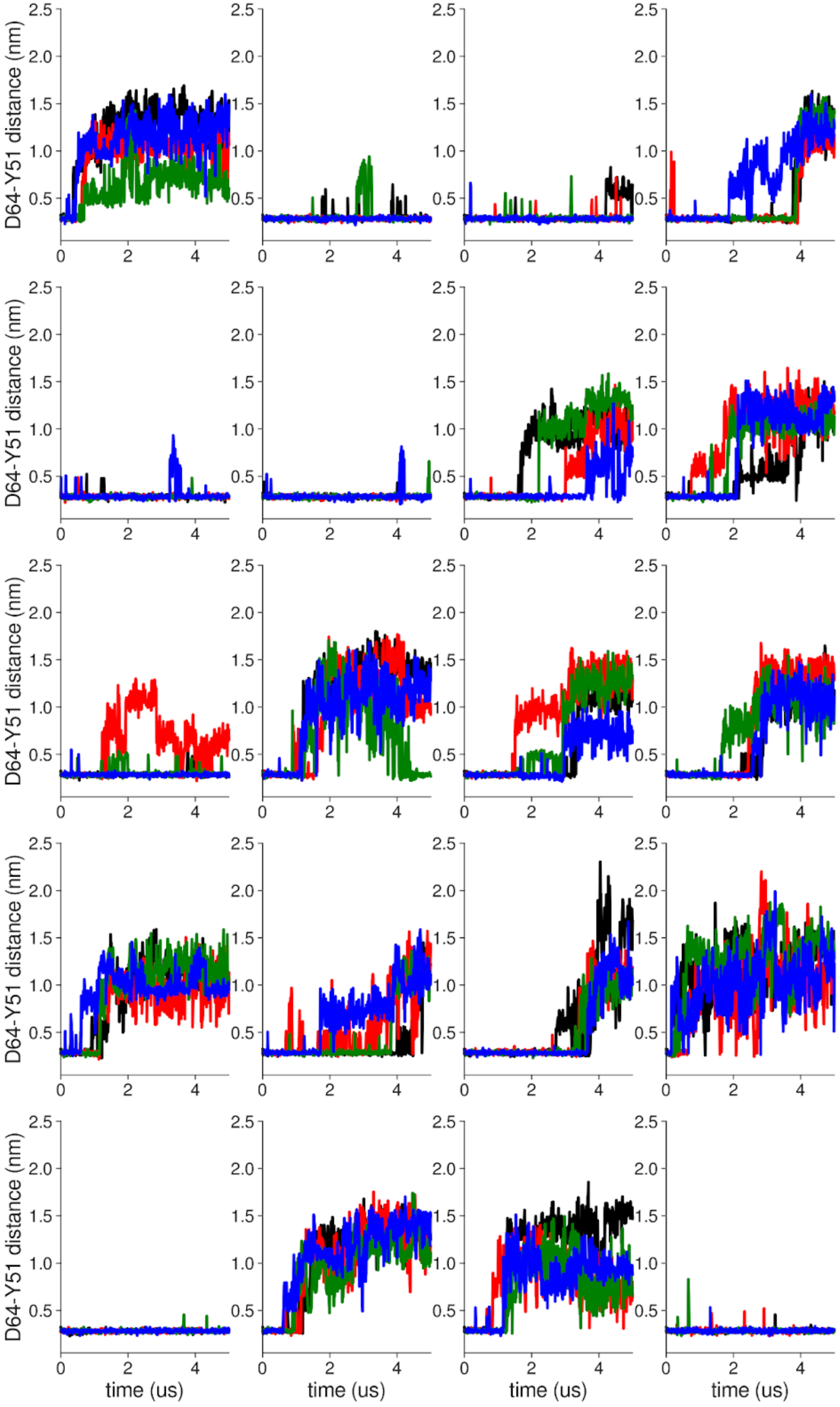


**Supplementary Figure 9 Individual distance traces between D64 CG atom and Y51 HH atom in long MthK V55E CHARMM simulations at 150 mV.** Each color refers to one of the channel monomers. Each panel shows traces from an independent, 5us long simulation. Distance values > 1 nm indicate broken interactions.

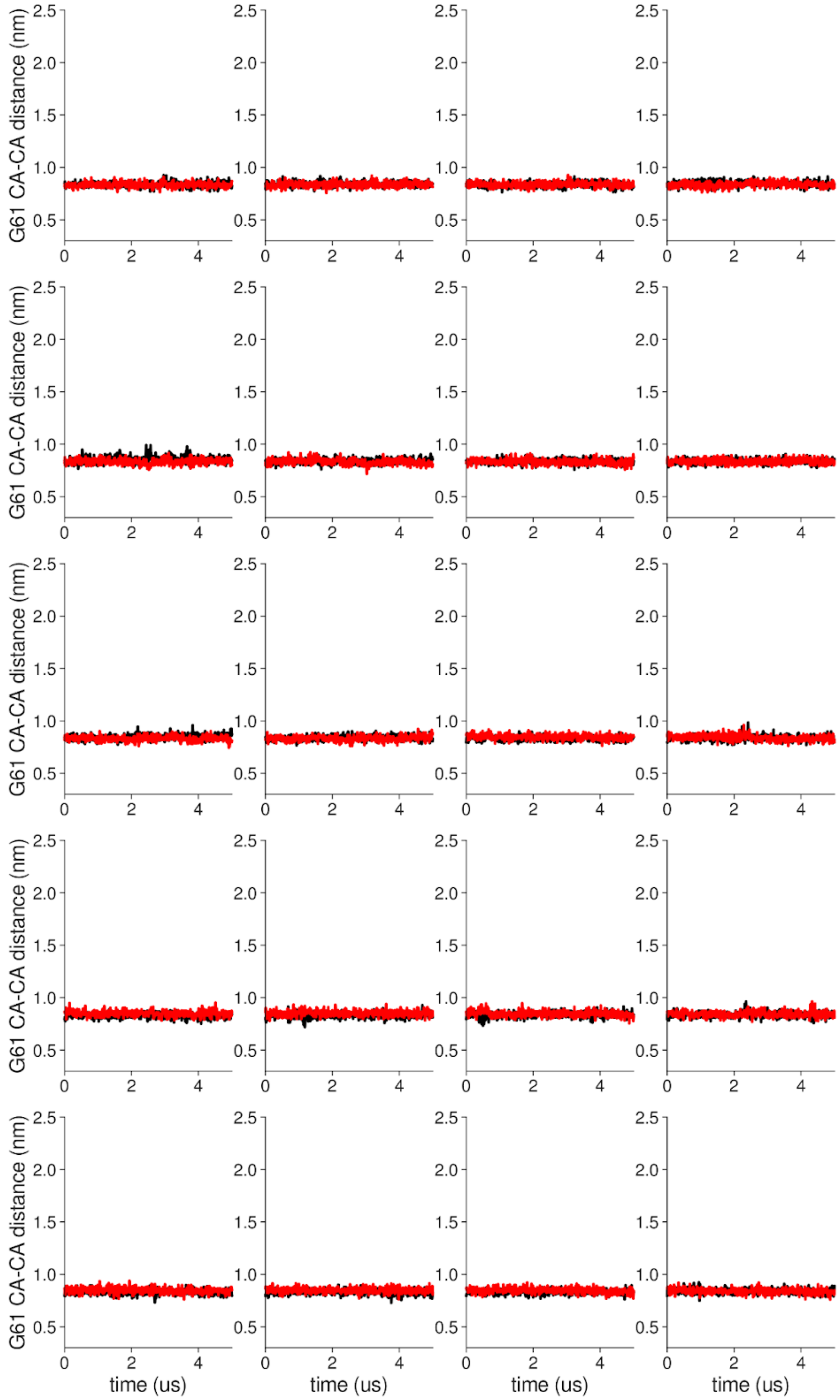


**Supplementary Figure 10 Individual distance traces between G61 CA atoms between oppositely oriented monomers in long MthK WT AMBER simulations at 300 mV.** Each panel shows traces from an independent, 5us long simulation.

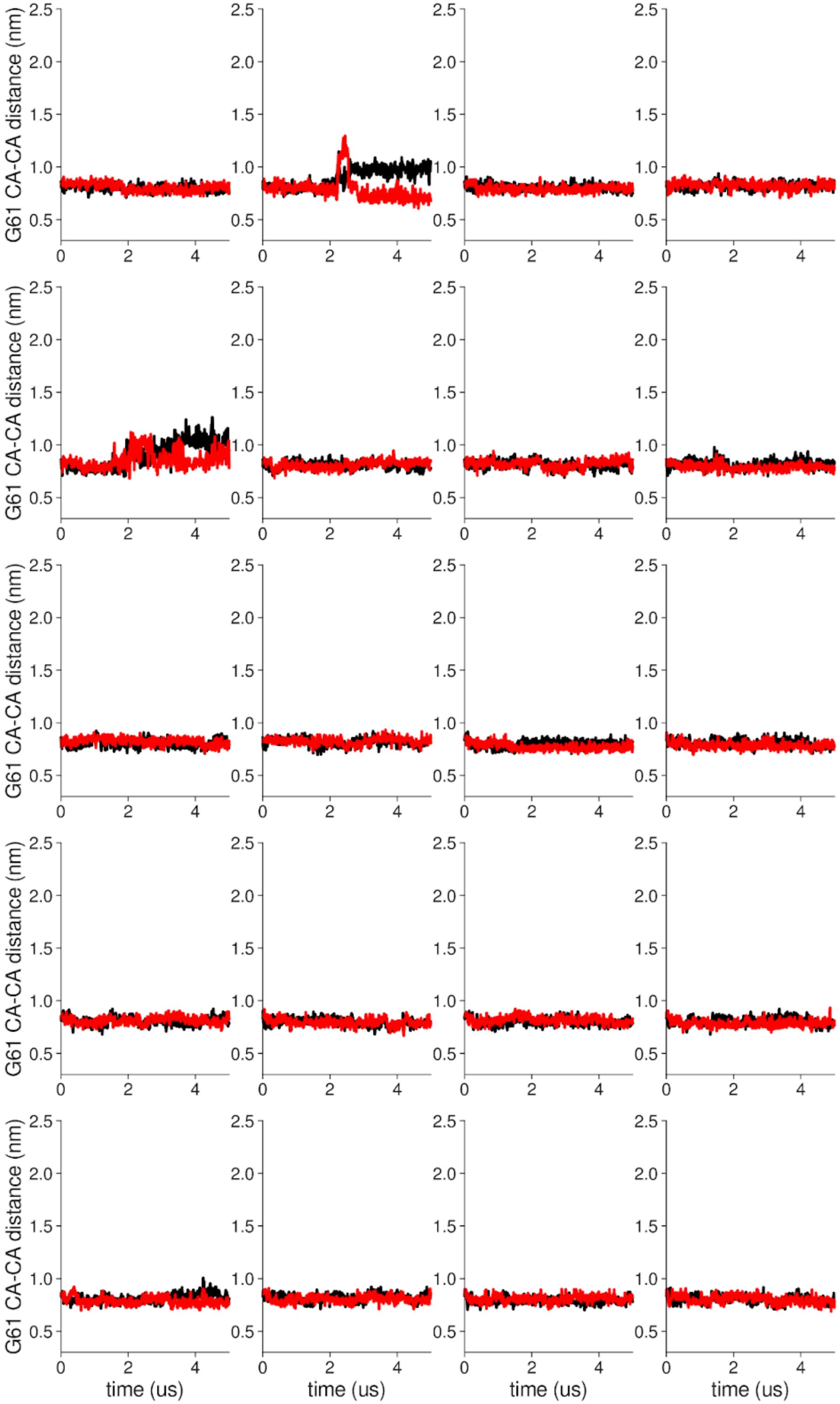


**Supplementary Figure 11 Individual distance traces between G61 CA atoms between oppositely oriented monomers in long MthK V55E AMBER simulations at 300 mV.** Each panel shows traces from an independent, 5us long simulation.

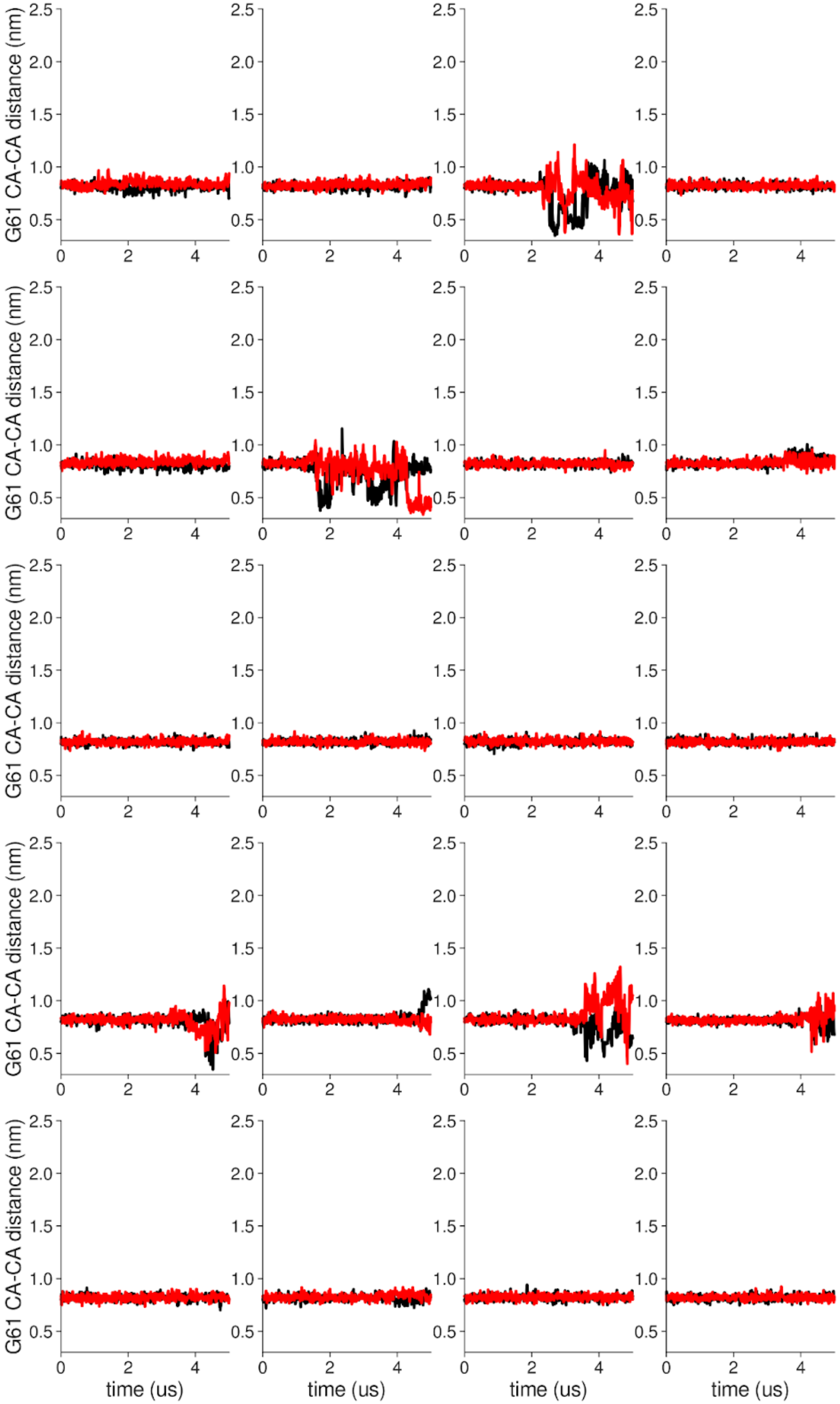


**Supplementary Figure 12 Individual distance traces between G61 CA atoms between oppositely oriented monomers in long MthK WT CHARMM simulations at 300 mV.** Each panel shows traces from an independent, 5us long simulation.

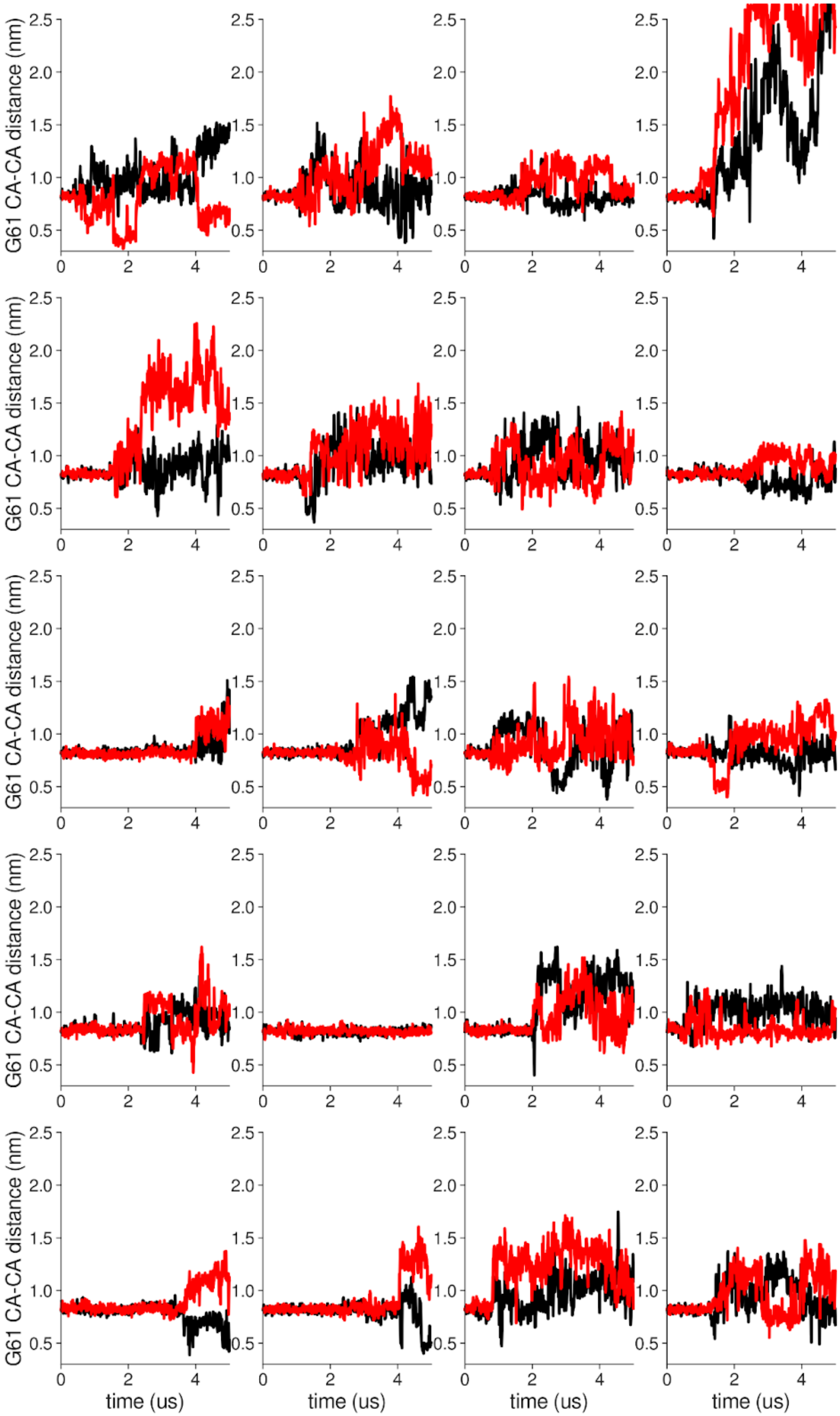


**Supplementary Figure 13 Individual distance traces between G61 CA atoms between oppositely oriented monomers in long MthK V55E CHARMM simulations at 300 mV.** Each panel shows traces from an independent, 5us long simulation.

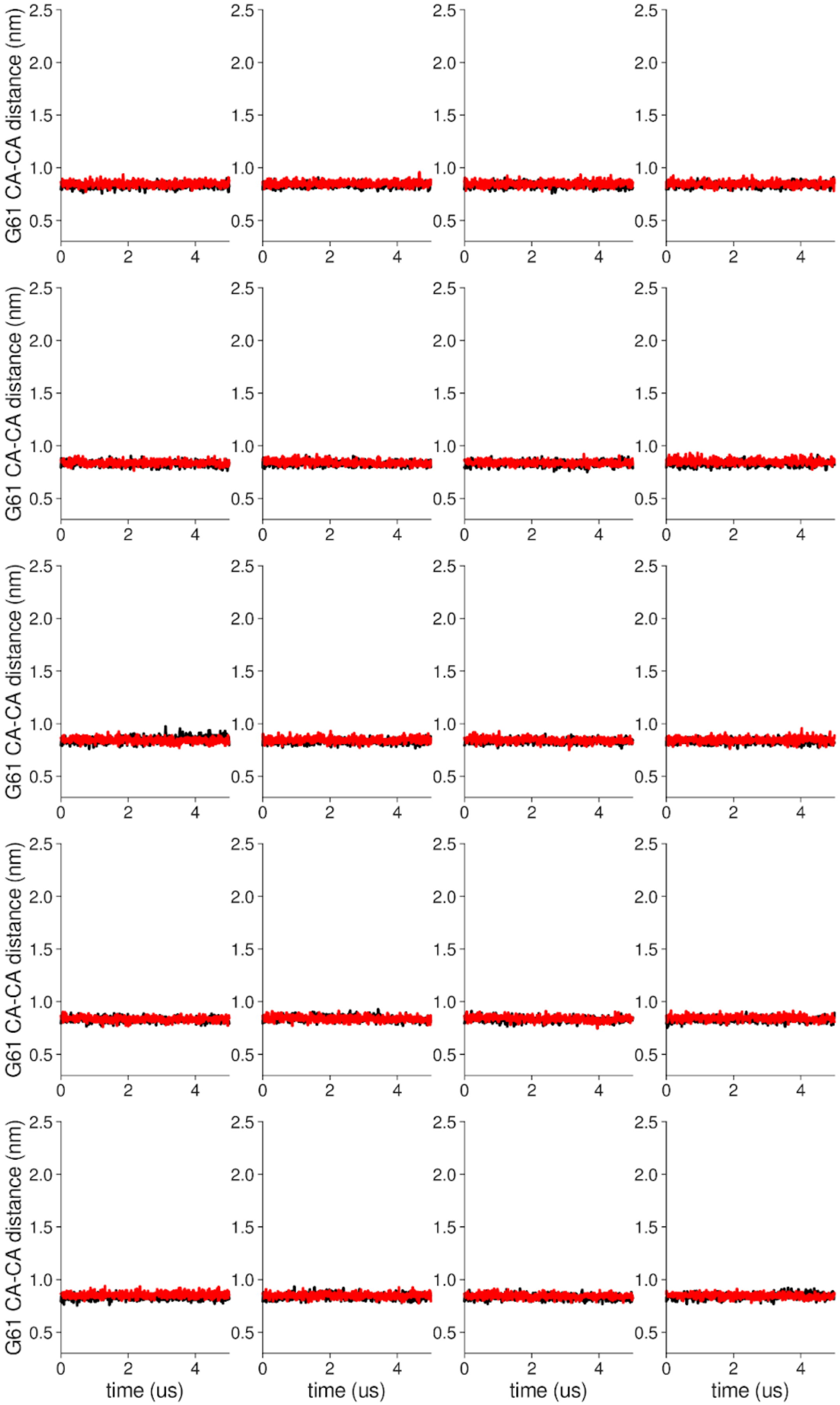


**Supplementary Figure 14 Individual distance traces between G61 CA atoms between oppositely oriented monomers in long MthK WT AMBER simulations at 150 mV.** Each panel shows traces from an independent, 5us long simulation.

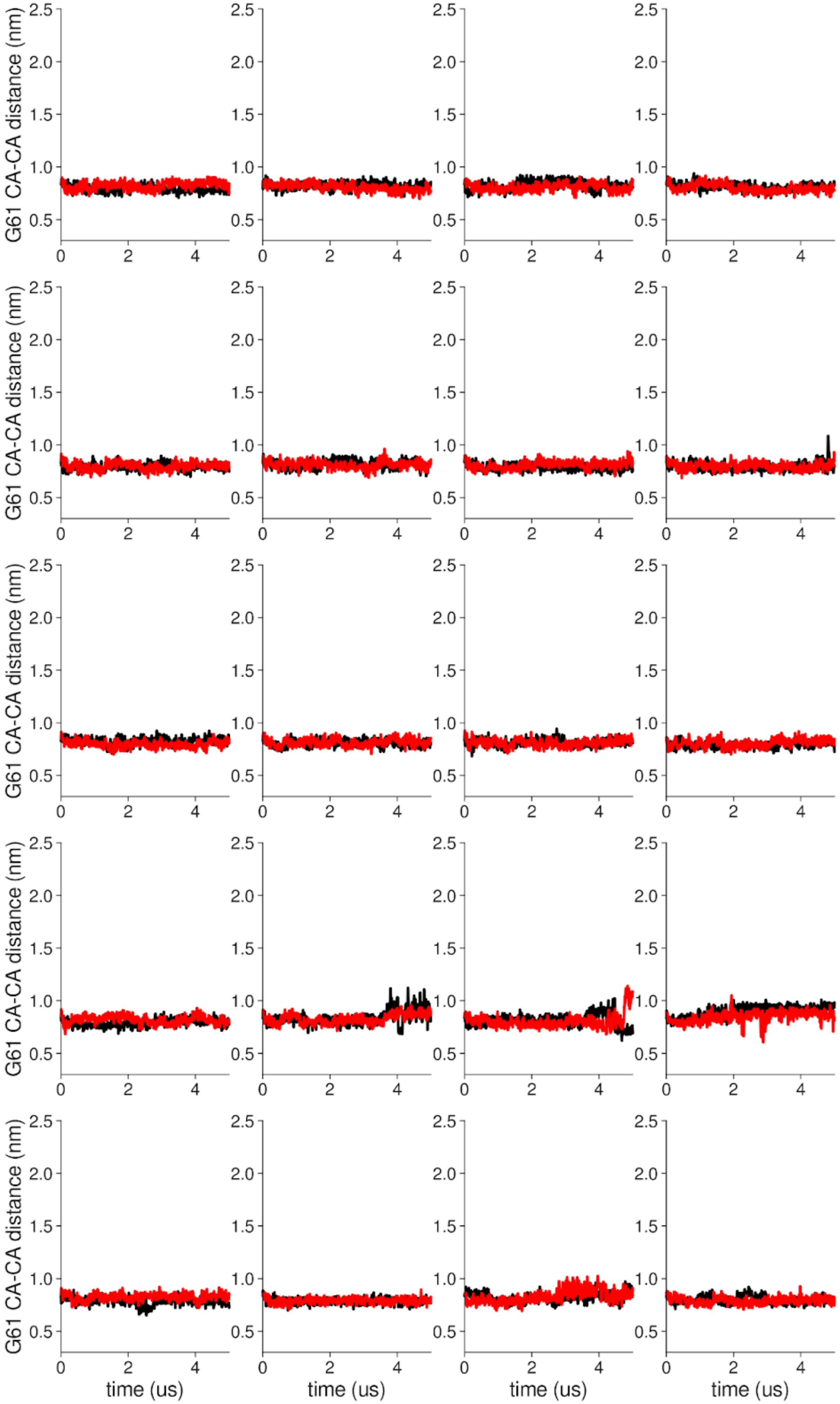


**Supplementary Figure 15 Individual distance traces between G61 CA atoms between oppositely oriented monomers in long MthK V55E AMBER simulations at 150 mV.** Each panel shows traces from an independent, 5us long simulation.

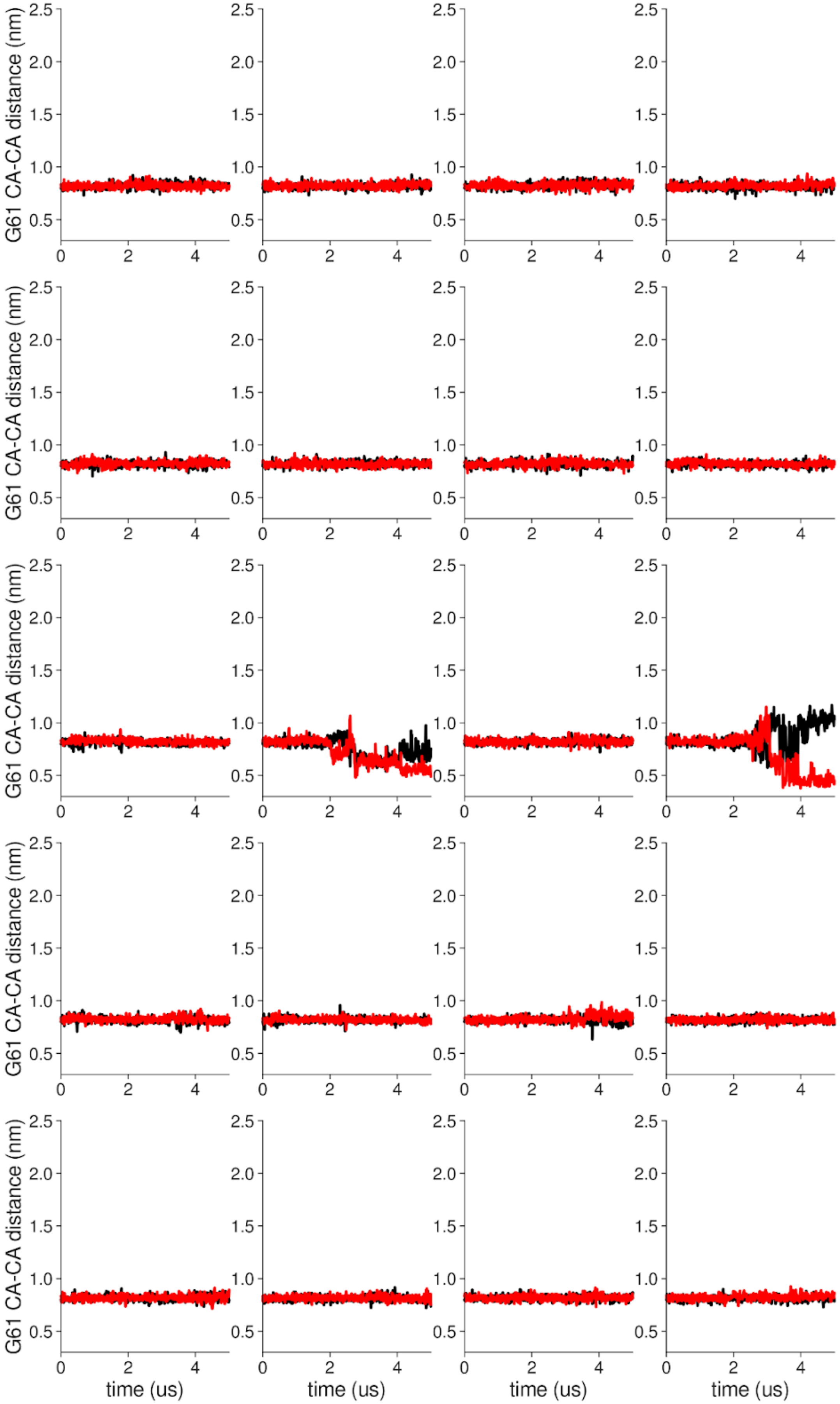


**Supplementary Figure 16 Individual distance traces between G61 CA atoms between oppositely oriented monomers in long MthK WT CHARMM simulations at 150 mV.** Each panel shows traces from an independent, 5us long simulation.

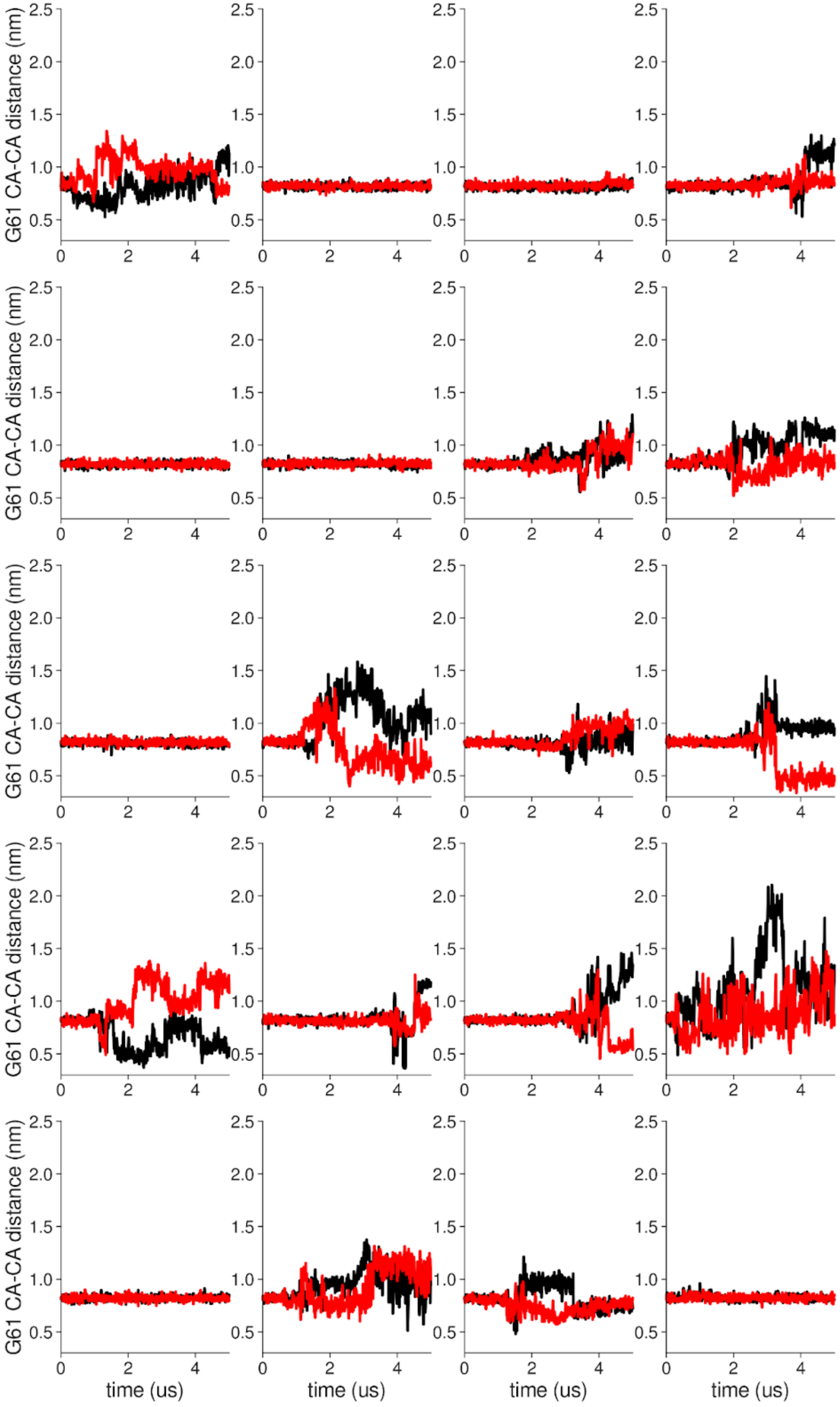


**Supplementary Figure 17 Individual distance traces between G61 CA atoms between oppositely oriented monomers in long MthK V55E CHARMM simulations at 300 mV.** Each panel shows traces from an independent, 5us long simulation.

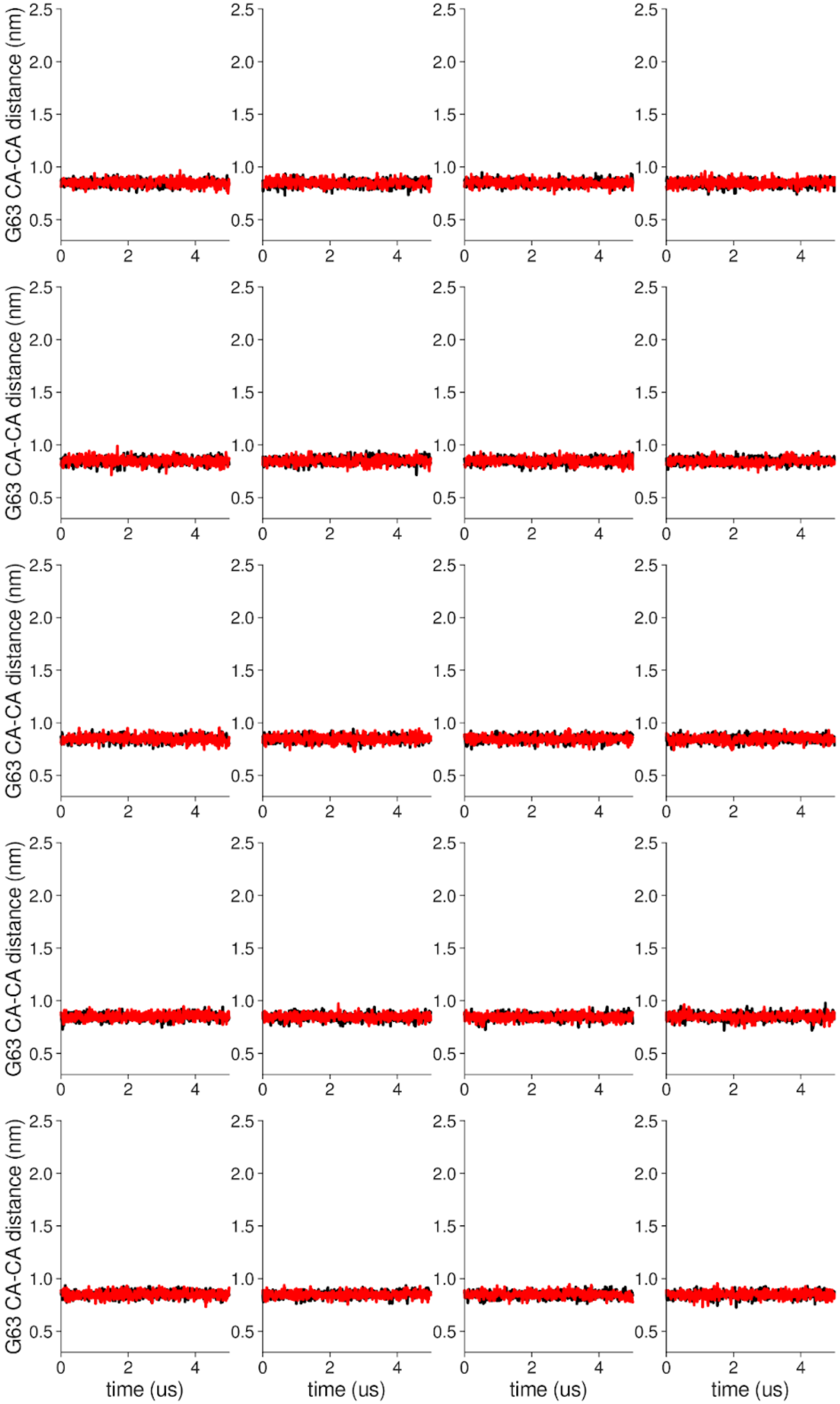


**Supplementary Figure 18 Individual distance traces between G63 CA atoms between oppositely oriented monomers in long MthK WT AMBER simulations at 300 mV.** Each panel shows traces from an independent, 5us long simulation.

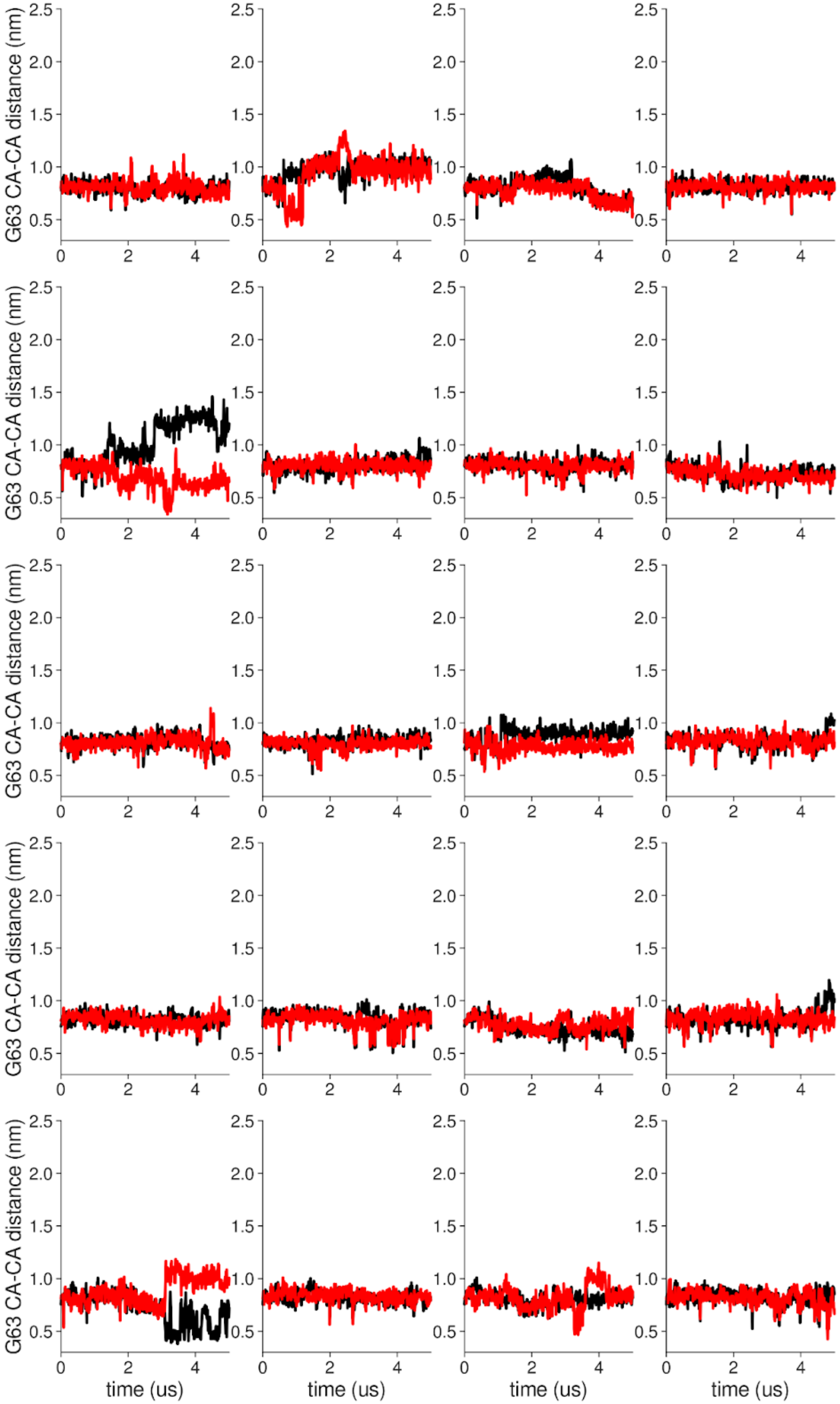


**Supplementary Figure 19 Individual distance traces between G63 CA atoms between oppositely oriented monomers in long MthK V55E AMBER simulations at 300 mV.** Each panel shows traces from an independent, 5us long simulation.

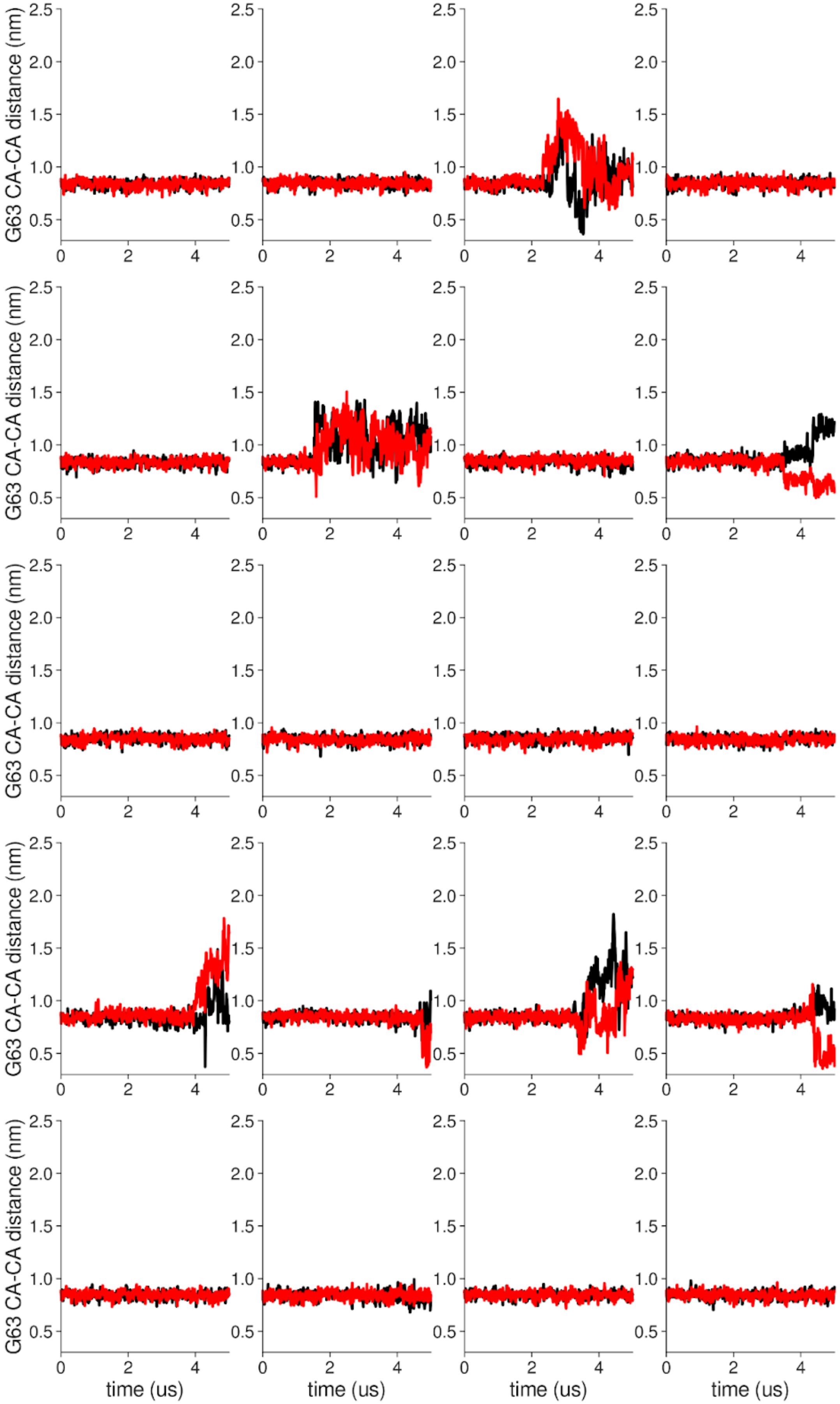


**Supplementary Figure 20 Individual distance traces between G63 CA atoms between oppositely oriented monomers in long MthK WT CHARMM simulations at 300 mV.** Each panel shows traces from an independent, 5us long simulation.

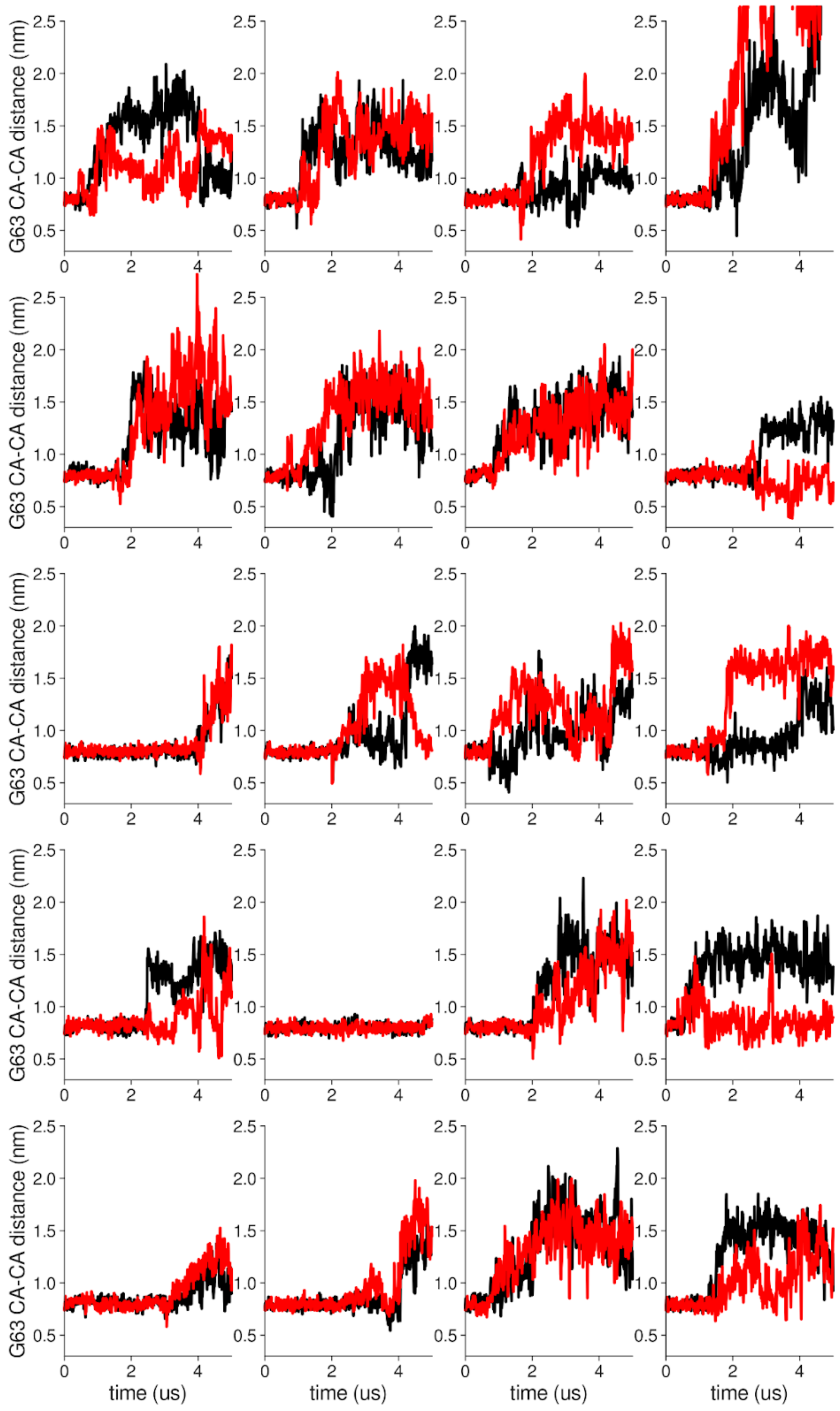


**Supplementary Figure 21 Individual distance traces between G63 CA atoms between oppositely oriented monomers in long MthK V55E CHARMM simulations at 300 mV.** Each panel shows traces from an independent, 5us long simulation.

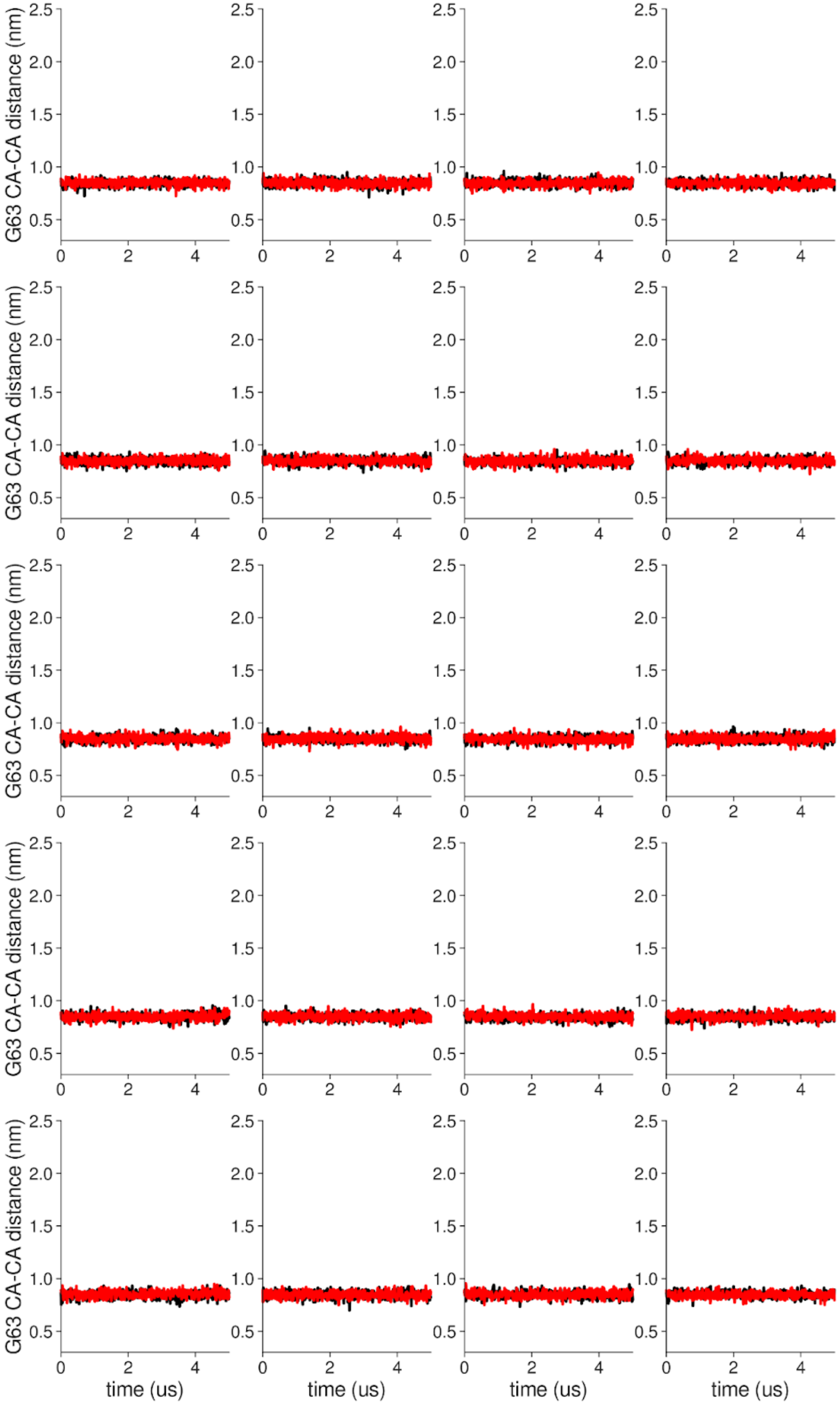


**Supplementary Figure 22 Individual distance traces between G63 CA atoms between oppositely oriented monomers in long MthK WT AMBER simulations at 150 mV.** Each panel shows traces from an independent, 5us long simulation.

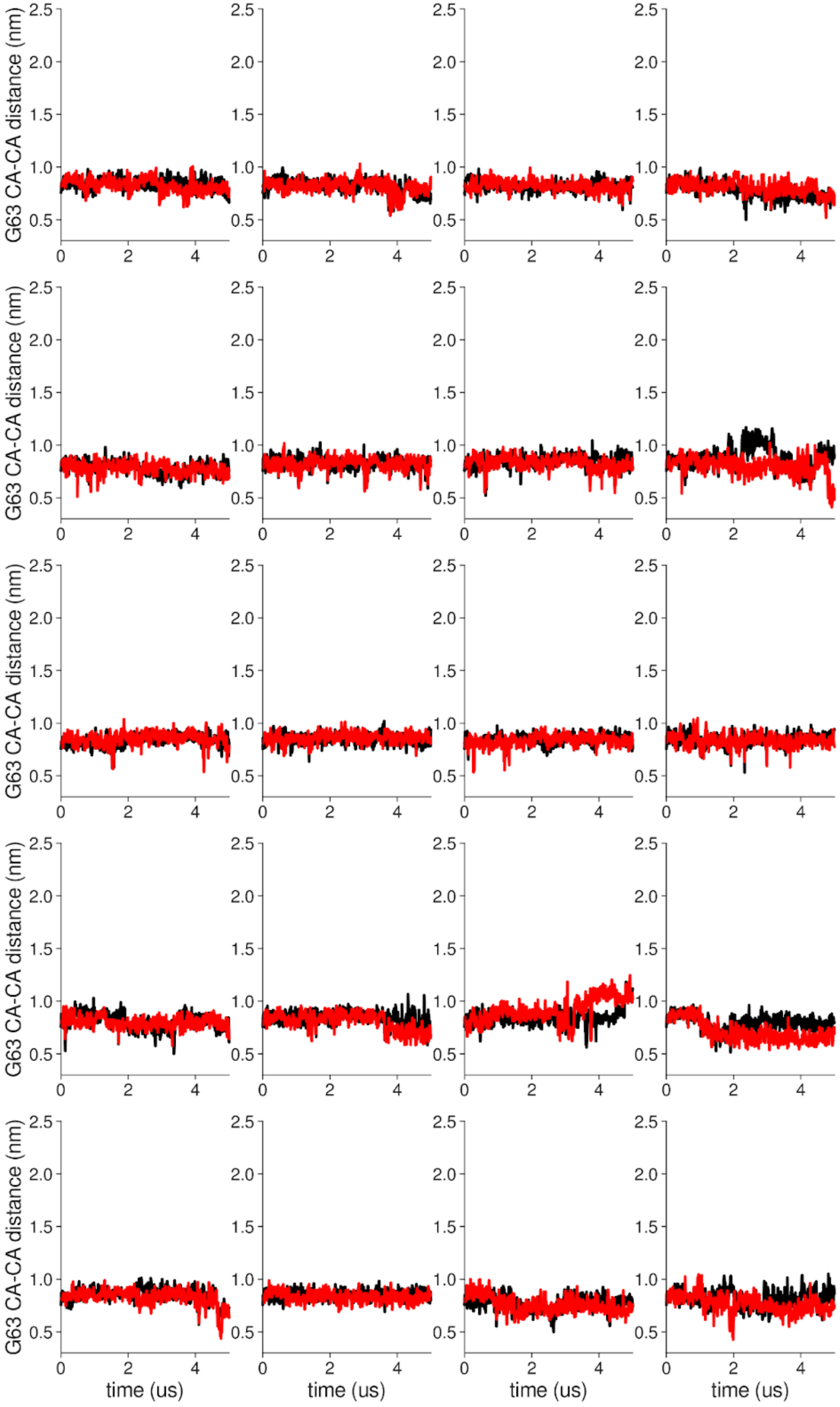


**Supplementary Figure 23 Individual distance traces between G63 CA atoms between oppositely oriented monomers in long MthK V55E AMBER simulations at 150 mV.** Each panel shows traces from an independent, 5us long simulation.

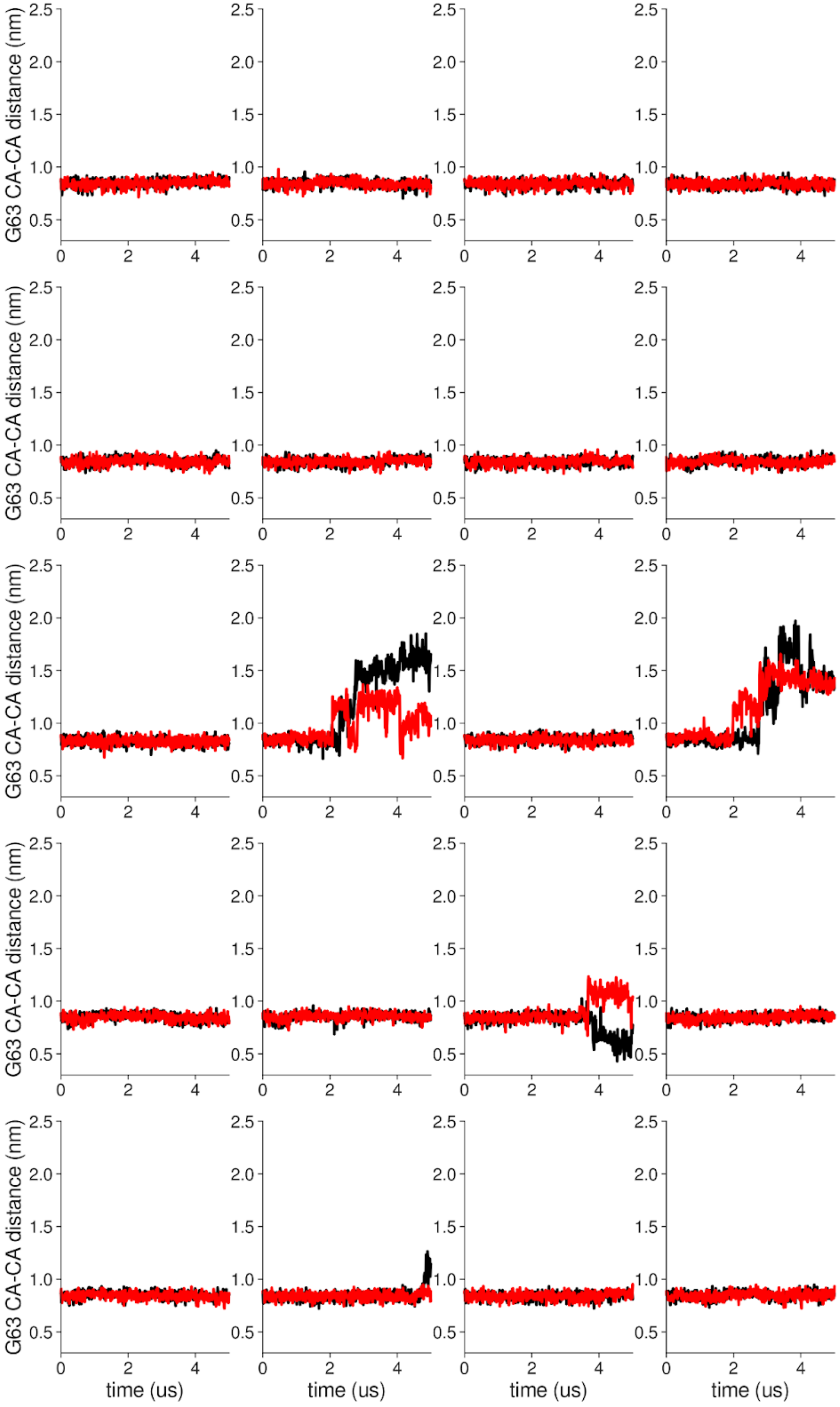


**Supplementary Figure 24 Individual distance traces between G63 CA atoms between oppositely oriented monomers in long MthK WT CHARMM simulations at 150 mV.** Each panel shows traces from an independent, 5us long simulation.

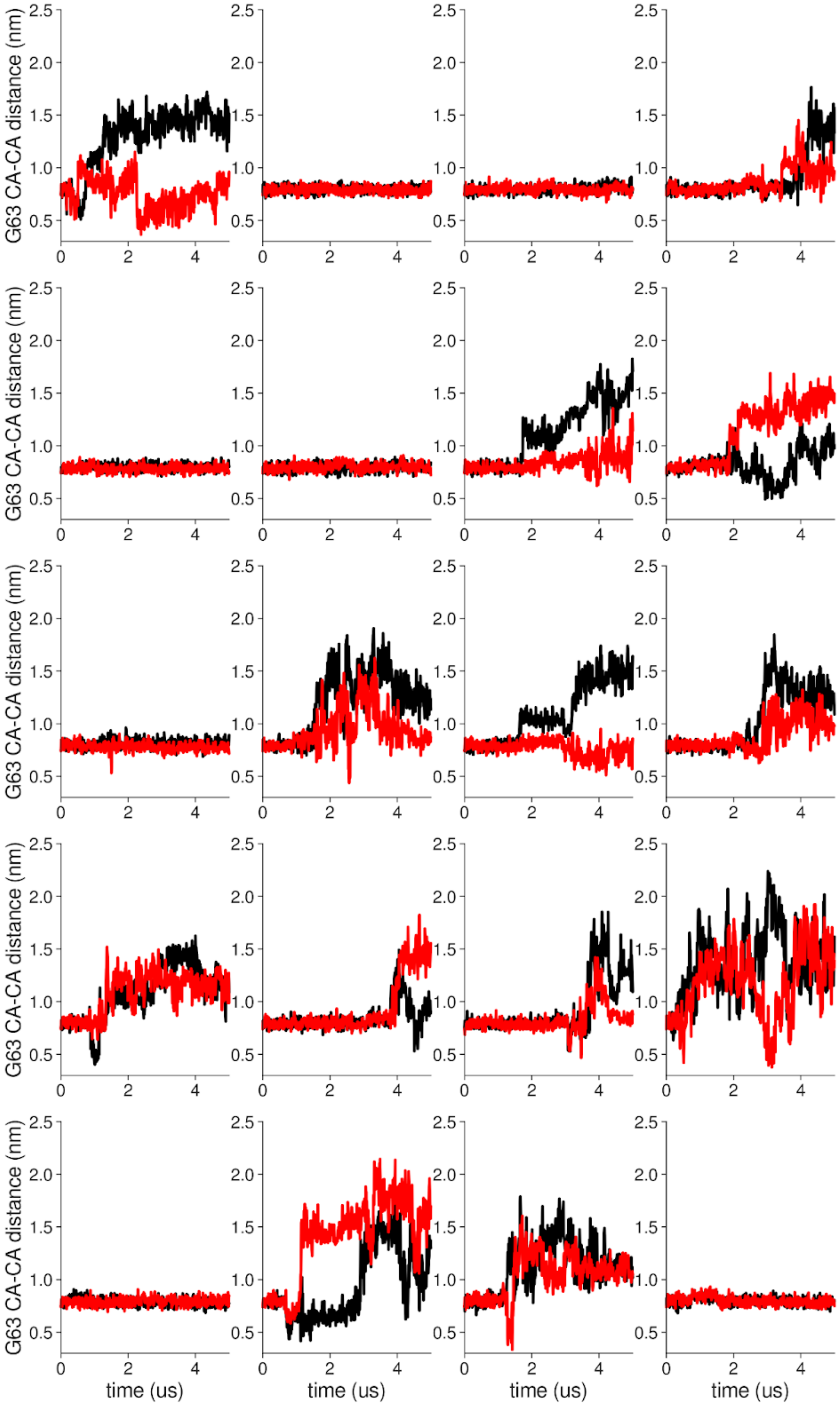


**Supplementary Figure 25 Individual distance traces between G63 CA atoms between oppositely oriented monomers in long MthK V55E CHARMM simulations at 150 mV.** Each panel shows traces from an independent, 5us long simulation.

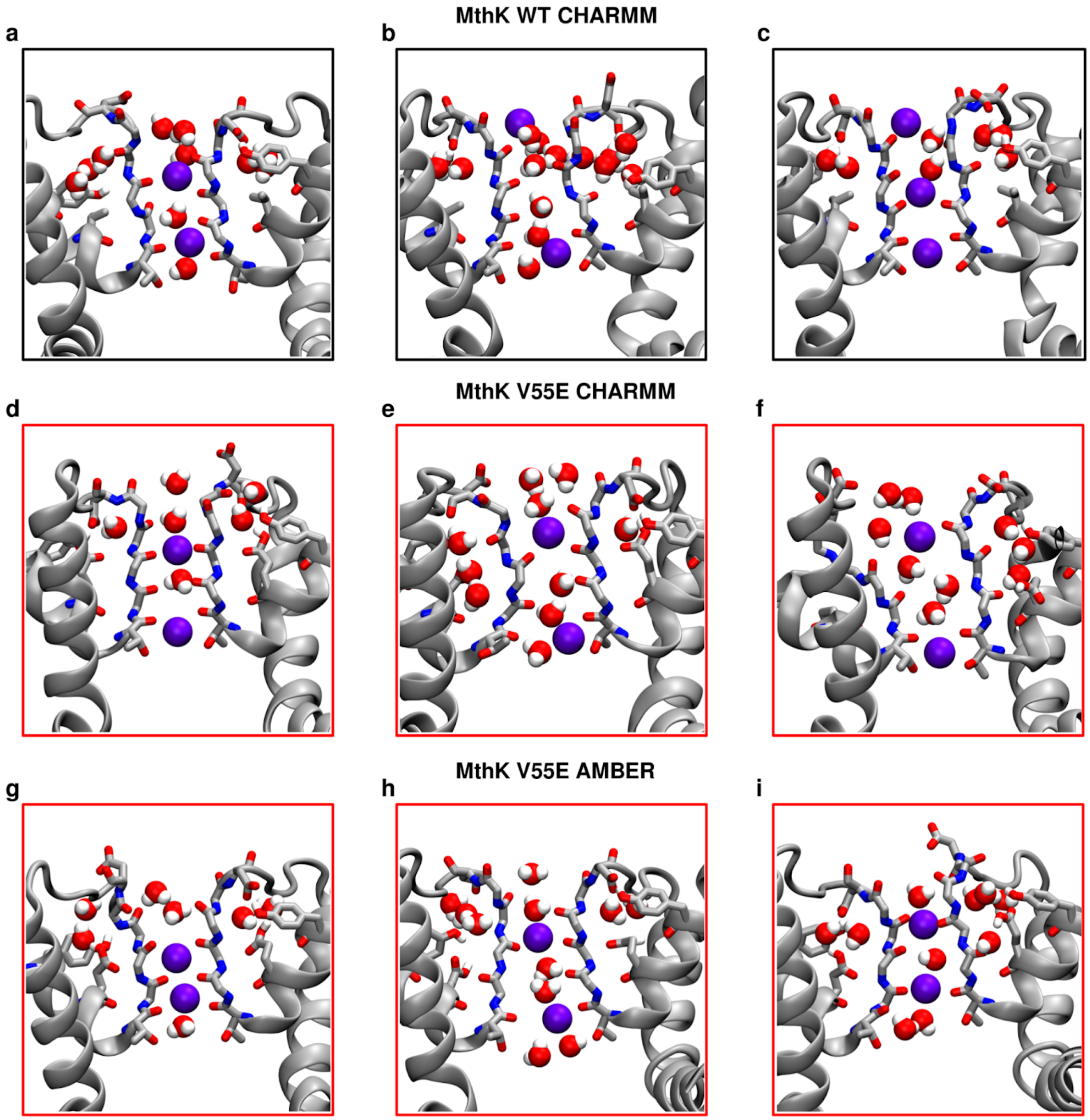


**Supplementary Figure 26 Conformations of inactivated filters in MD simulations of MthK WT and MthK V55E at 150 mV.** A-C, Snapshots from simulations of MthK WT with the CHARMM force field, D-F, Snapshots from simulations of MthK V55E with the CHARMM force field, G-I, Snapshots from simulations of MthK V55E with the AMBER force field. Both aspartate (D64) sidechain flipping as well as valine (V60) and glycine (G61) carbonyl flipping can be observed when SFs lose potassium ions.

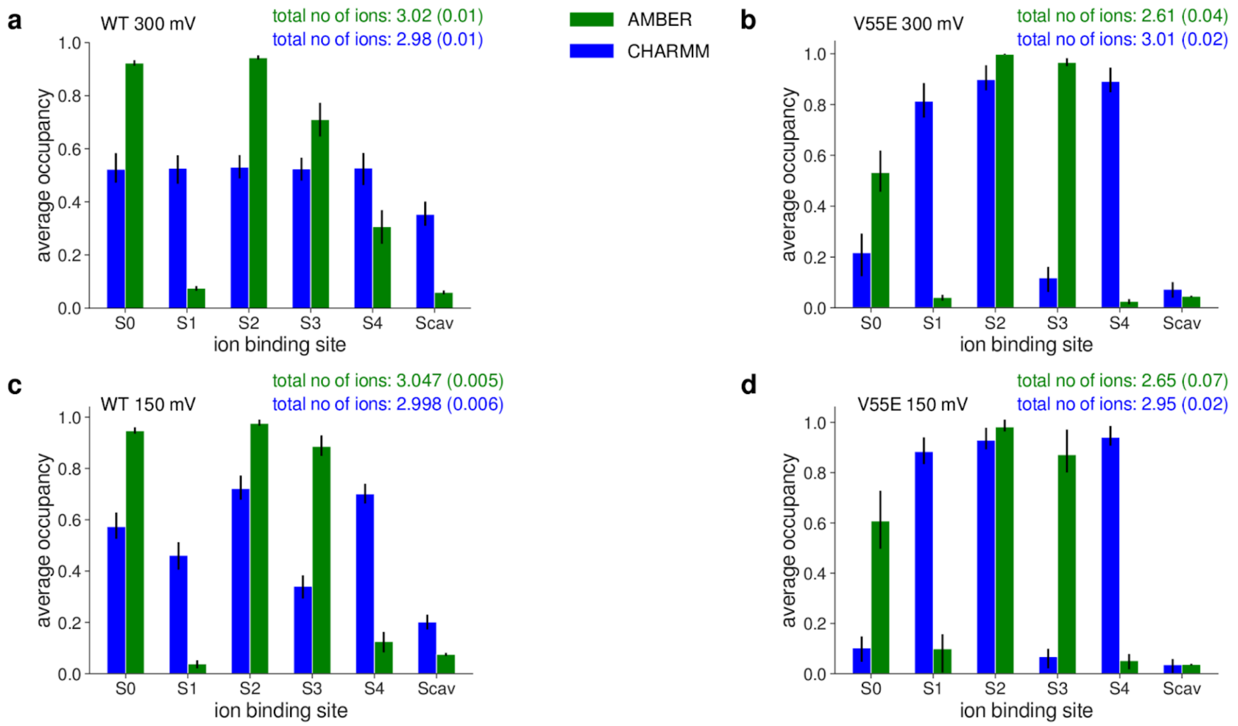


**Supplementary Figure 27 Average potassium occupancy of each ion binding site in the SF of MthK WT and V55E before the D64 flip.** A, MthK WT at 300 mV, B, MthK V55E at 300 mV, C, MthK WT at 150 mV, D, MthK V55E at 150 mV.

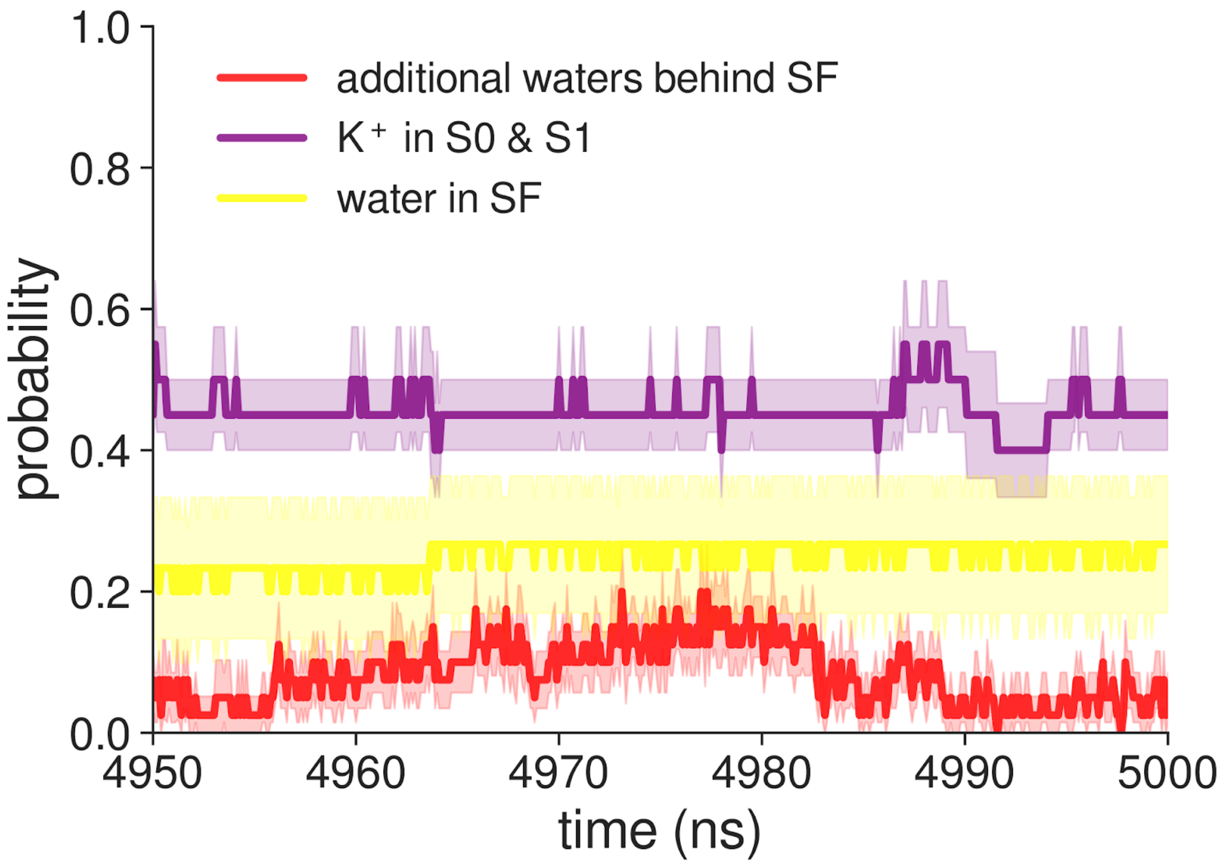


**Supplementary Figure 28 Probabilities of molecular events in the last 50ns of MD simulations of MthK WT with the AMBER force field.**

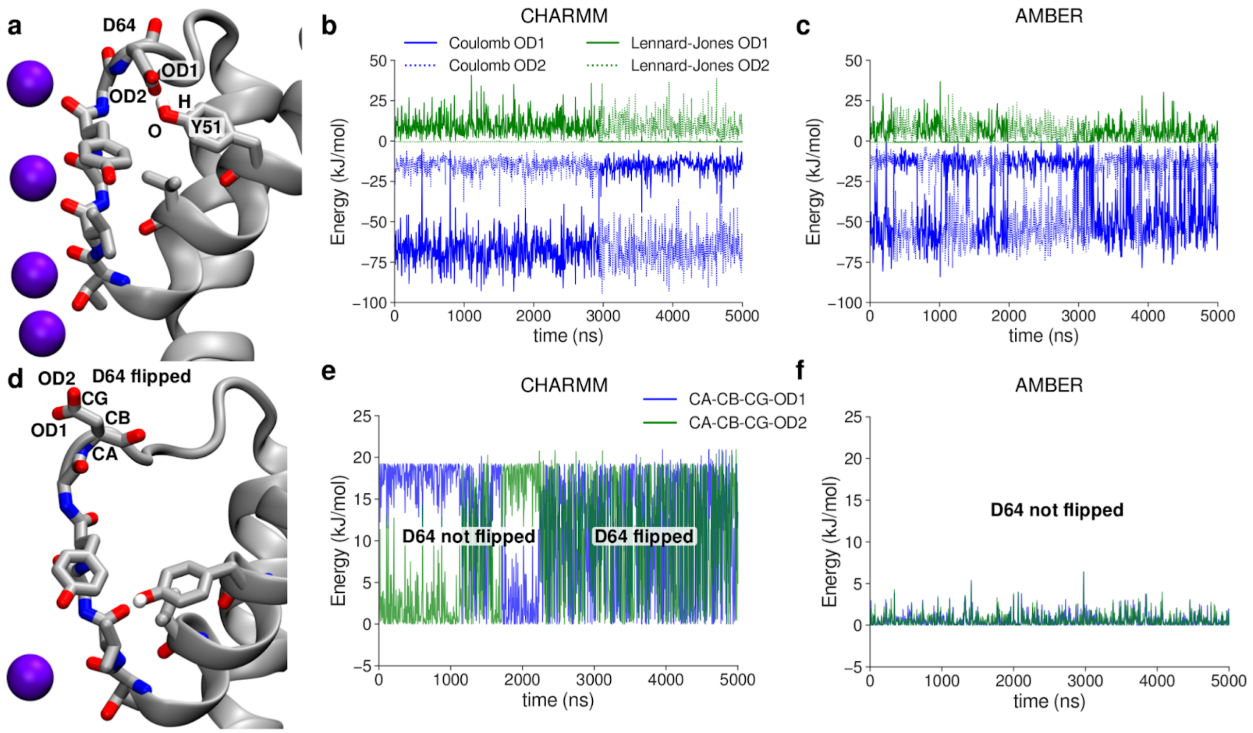


**Supplementary Figure 29 Factors determining the dynamics of the D64 side chain in MthK WT in CHARMM and AMBER force fields.** A, Typical position of side chains before the aspartate flip. The D64 side chain interacts with the Y51 side chain via hydrogen bond. B, C Interaction energies (Coulomb and Lennard-Jones) between atoms involved in D64-Y51 hydrogen bond, in CHARMM and AMBER, respectively. D, Snapshot after the aspartate flip, showing the D64 side chain pointing away from the protein. E, F, Energies of the two dihedrals angles defined by heavy atoms from the D64 side chain, in CHARMM and AMBER, respectively.

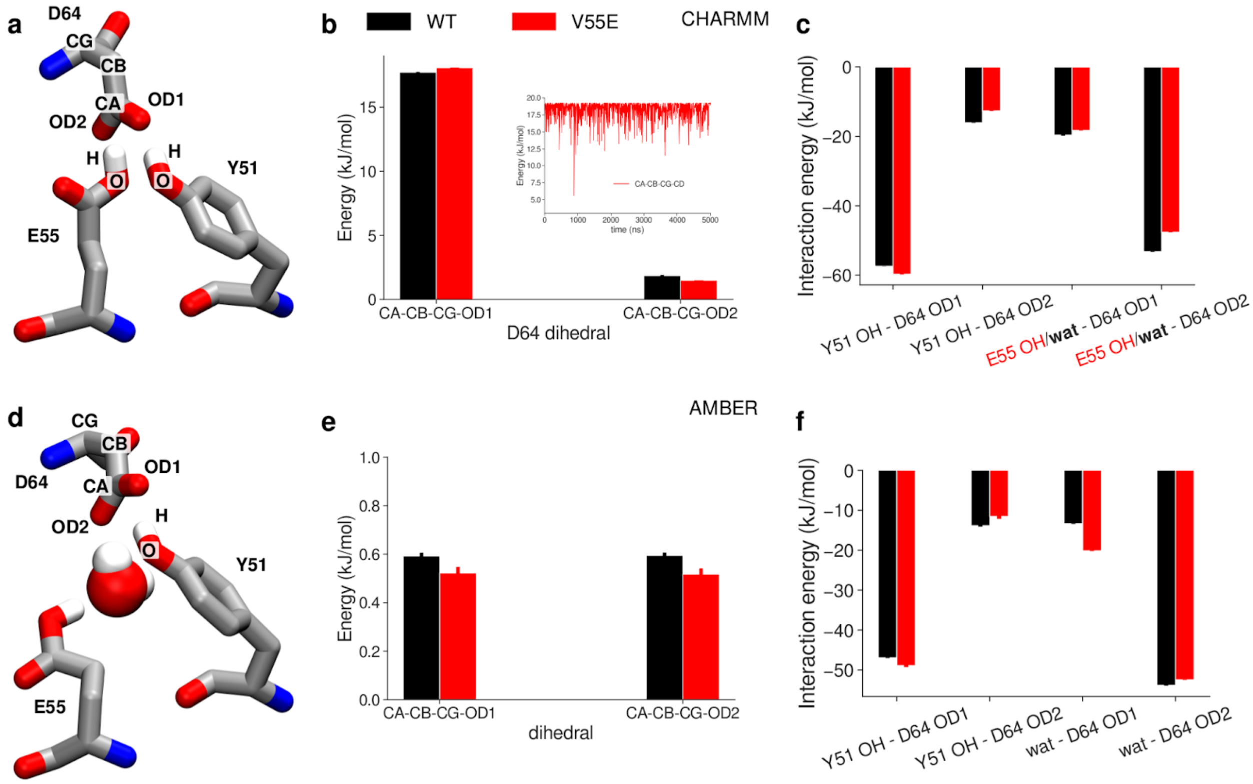


**Supplementary Figure 30 The effects of the V55E mutation on factors determining the dynamics of the D64 side chain in CHARMM and AMBER force fields.** A, Typical position of D64, E55 and Y51 side chains in the CHARMM force field, before the aspartate flip. The E55 side chain is in the 'vertical' orientation. B, Effects of the V55E mutation on the average energy of the two dihedrals angles of the D64 side chain. Inset shows a typical time evolution of a high energy dihedral. C, Average interaction energies between atoms forming the Y51-D64 hydrogen bond and E55-D64 hydrogen bond (red, in MthK V55E) or D64-water hydrogen bond (black, in MthK WT). D, Typical position of D64, E55 and Y51 side chain the AMBER force field, before the aspartate flip. The E55 side chain is in the 'horizontal' orientation, and there is an additional water molecule between D64 and E55 side chains. E, same as B, but in the AMBER force field. F, same as C, but in the AMBER force field. As the E55 side chain is typically in the horizontal orientation, and thus does not form a hydrogen bond with the D64 side chain, only interaction energies with water are included.

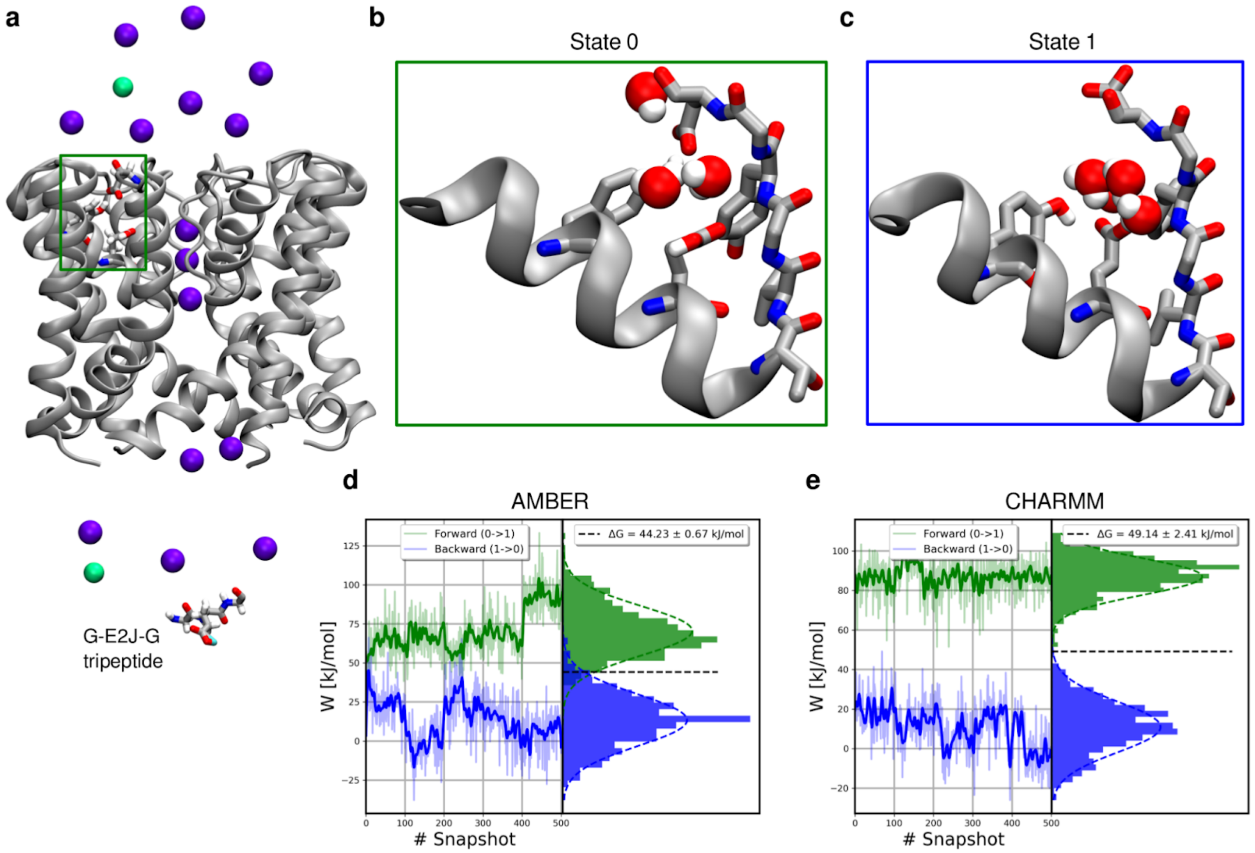


**Supplementary Figure 31 Nonequilibrium free energy calculations of E55 deprotonation.** A, System used in free energy calculations, showing a selected E55 glutamate (green box), a reference state tripeptide G-E2J-G and a lowered salt concentration of 150 mM. B, C, End states for E55 in free energy simulations: in the State 0, E55 is protonated and in the State 1 it is deprotonated. In both states, E55 is kept in its horizontal orientation. When E55 is deprotonated (State 1), the side chain of D64 can flip toward the extracellular space. D, E, Distributions of work values in nonequilibrium transitions and the final estimates of $\Delta G$ obtained with pmx, in AMBER and CHARMM, respectively.

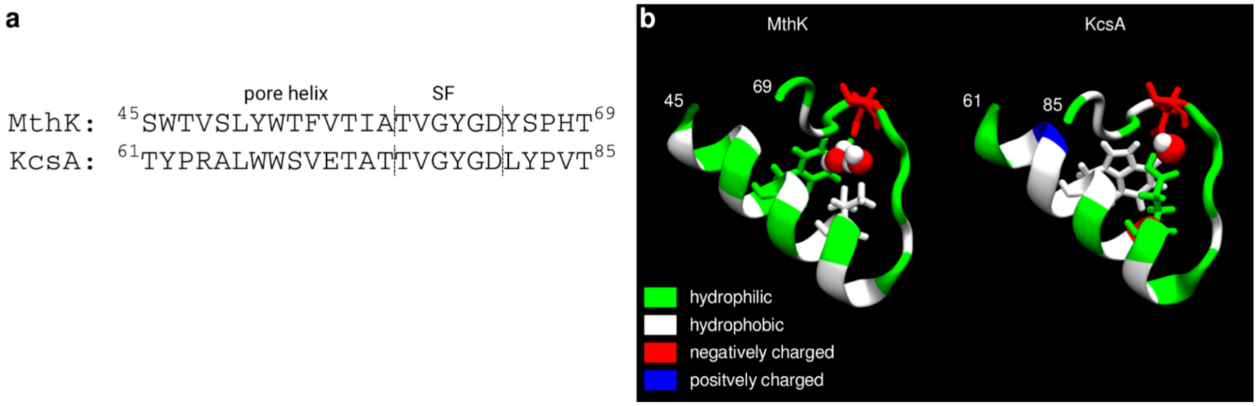


**Supplementary Figure 32 Comparison of the pore region between MthK and KcsA channels.** A, Sequence alignment for MthK and KcsA. B, Visualization of the pore helix, selectivity filter and the loop following SF in MthK and KcsA. Residues are colored according to their chemical character: hydrophilic (green), hydrophobic (white), negatively charged (red) or positively charged (blue). The critical residues discussed in this work: Y51/W67, V55/E71 and D64/D80 (in MthK/KcsA) are shown as sticks. Water molecules are shown as red and white spheres.
